# Simulation of closed-loop deep brain stimulation control schemes for suppression of pathological beta oscillations in Parkinson’s disease

**DOI:** 10.1101/2019.12.26.880781

**Authors:** John E. Fleming, Eleanor Dunn, Madeleine M. Lowery

**Author notes:** Correspondence: John Fleming.

## Abstract

This study presents a computational model of closed-loop control of deep brain stimulation (DBS) for Parkinson’s disease (PD) to investigate clinically-viable control schemes for suppressing pathological beta-band activity. Closed-loop DBS for PD has shown promising results in preliminary clinical studies and offers the potential to achieve better control of patient symptoms and side effects with lower power consumption than conventional open-loop DBS. However, extensive testing of algorithms in patients is difficult. The model presented provides a means to explore a range of control algorithms *in silico* and optimize control parameters before preclinical testing. The model incorporates (i) the extracellular DBS electric field, (ii) antidromic and orthodromic activation of STN afferent fibers, (iii) the LFP detected at non-stimulating contacts on the DBS electrode and (iv) temporal variation of network beta-band activity within the thalamo-cortico-basal ganglia loop. The performance of on-off and dual-threshold controllers for suppressing beta-band activity by modulating the DBS amplitude were first verified, showing levels of beta suppression and reductions in power consumption comparable with previous clinical studies. Proportional (P) and proportional-integral (PI) closed-loop controllers for amplitude and frequency modulation were then investigated. A simple tuning rule was derived for selecting effective PI controller parameters to target long duration beta bursts while respecting clinical constraints that limit the rate of change of stimulation parameters. Of the controllers tested, PI controllers displayed superior performance for regulating network beta-band activity whilst accounting for clinical considerations. Proportional controllers resulted in undesirable rapid fluctuations of the DBS parameters which may exceed clinically tolerable rate limits. Overall, the PI controller for modulating DBS frequency performed best, reducing the mean error by 83% compared to DBS off and the mean power consumed to 25% of that utilized by open-loop DBS. The network model presented captures sufficient physiological detail to act as a surrogate for preclinical testing of closed-loop DBS algorithms using a clinically accessible biomarker, providing a first step for deriving and testing novel, clinically-suitable closed-loop DBS controllers.

## 1 Introduction

In recent years, there has been growing interest among the clinical and scientific communities on the potential offered by ‘closed-loop’ DBS. In a closed-loop DBS configuration, the patient’s clinical state is quantified and utilized to alter stimulation parameters as necessary, so the required stimulation to minimize their disease symptoms is delivered, thus reducing potential stimulation induced side-effects while controlling symptoms. A critical step in the development of such systems is the identification of signal features or ‘biomarkers’ which have the potential to quantify the clinical state. One of the most promising features examined for closed-loop control of DBS in PD is the level of beta-band (10 – 30 Hz) oscillatory activity within the subthalamic nucleus (STN) and cortico-basal ganglia network. Pathological exaggerated activity within this frequency band is correlated with motor impairment and its suppression, due to medication or DBS, with motor improvement (Kühn et al., 2008, 2009; Silberstein et al., 2005b, 2005a). This oscillatory activity, however, is not continuously elevated, but rather fluctuates between long, greater than 400 ms, and short duration bursts of beta activity, with only long burst durations being positively correlated with motor impairment in PD (Tinkhauser et al., 2017b, 2017a). These features, in combination with the potential to record LFP activity during stimulation from non-stimulating contacts on the DBS electrode, render it an appealing biomarker for closed-loop DBS.

Closed-loop DBS for PD utilizing LFP derived measures of beta-band oscillatory activity has been successfully tested in small cohorts of PD patients over relatively short timescales. These studies have examined amplitude modulation of the DBS waveform in response to changes in the LFP beta-band activity. ‘On-off’ stimulation strategies, where DBS is triggered on or off as the measured oscillatory activity crosses a desired threshold value, were the first closed-loop strategies tested in patients (Little et al., 2013, 2016). Although these strategies offer benefits with respect to traditional open-loop stimulation, they rely on optimal stimulation parameters that are identified during open-loop, continuous DBS. If these stimulation parameters are no longer effective, for example, due to diurnal changes in beta activity, variations in the electrode impedance, or as the disease progresses, the controller is unable to adapt and delivers suboptimal performance. Velisar *et al*. proposed an alternative ‘dual-threshold’ algorithm where the amplitude of the DBS waveform is systematically increased, decreased or kept constant as the measured LFP beta-band activity remains above, below or within a desired target range (Velisar et al., 2019). Although the strategy can maintain the beta activity within a target range, it remains a relatively simple form of control where the DBS amplitude is varied at a fixed rate if the beta-band activity lies outside the target range. Arlotti *et al.* and Rosa *et al.* investigated an alternative approach where the DBS waveform was linearly modulated in response to the measured LFP beta-band activity in freely moving PD patients (Arlotti et al., 2018; Rosa et al., 2015). Proportional amplitude modulation stimulation strategies such as this, where the DBS amplitude is varied proportionally to the measured LFP beta-band activity, potentially offer more benefits than on-off and dual-threshold strategies, in theory, because they ideally only deliver the stimulation required to reduce beta-band LFP activity to suppress PD symptoms.

In conjunction with the amplitude modulation stimulation strategies that have been investigated so far, control theory offers a wealth of control schemes which may potentially offer better control of patient symptoms and side effects, whilst minimizing battery consumption, over the current state-of-the-art strategies. The development of novel, effective control schemes for DBS, however, is challenging and trialing in humans or animals is difficult due to its invasive nature. Computational modelling provides an alternative approach for designing and testing more complex forms of closed-loop DBS control. Although computational models have been previously used to investigate closed-loop control strategies for DBS (Gorzelic et al., 2013; Grant and Lowery, 2013; Haidar et al., 2016; Liu et al., 2017; Popovych and Tass, 2019; Santaniello et al., 2011; Su et al., 2019), they typically do not relate well to clinically relevant parameters and, in particular, rarely incorporate both simulation of the LFP and extracellular application of DBS. Simulation of the LFP is desirable for developing computational models that can be readily translated to patients as the LFP is currently the most accessible biomarker for closed-loop DBS in PD (Priori et al., 2013). In addition, simulation of the electric field and extracellular application of DBS to axons and branching afferents is necessary to enable variations in DBS amplitude to be simulated. To bridge the link between computational approaches and clinically-viable closed-loop approaches it is thus necessary to develop a model which captures the dynamics of the relevant neural system, the electric field generated by DBS, and the resulting LFP recording.

To address this, the aim of this study was to develop a physiologically based model of the cortico-basal ganglia network, which incorporates extracellular DBS and simulation of the STN LFP, that can be utilized to test clinically relevant closed-loop DBS control strategies. The developed model captures increased network beta-band oscillatory activity and simulates the synaptically generated STN LFP and extracellular application of DBS to STN afferent fibers, including antidromic activation of cortical pathways. The performance of on-off and dual-threshold amplitude modulation strategies are verified in the model before the feasibility of proportional (P) and proportional-integral (PI) control schemes for modulating DBS amplitude or frequency are investigated. The model provides an *in silico* testbed for developing new closed-loop DBS control strategies using STN LFP-derived features in PD.

## 2 Materials and Methods

The structure of the network model of DBS is presented in Figure 1 and includes the closed loop formed between the cortex, basal ganglia and thalamus (Nambu et al., 2002; Parent and Hazrati, 1995). The model extends previous network models of the parkinsonian cortico-basal ganglia during DBS (Hahn and McIntyre, 2010; Kang and Lowery, 2013, 2014; Kumaravelu et al., 2016; Rubin and Terman, 2004; Terman et al., 2002) by (i) incorporating the extracellular DBS electric field (ii) simulating antidromic and orthodromic activation of cortical and globus pallidus efferent fibers to the STN and (iii) simulating the LFP detected at non-stimulating contacts on the DBS electrode due to STN synaptic activity and (iv) mimics temporal variation of network beta activity within the thalamo-cortico-basal ganglia loop. The model was used to investigate the performance of closed-loop amplitude and frequency modulation strategies using an LFP derived measure of the network beta-band oscillatory activity.

**Figure 1:**
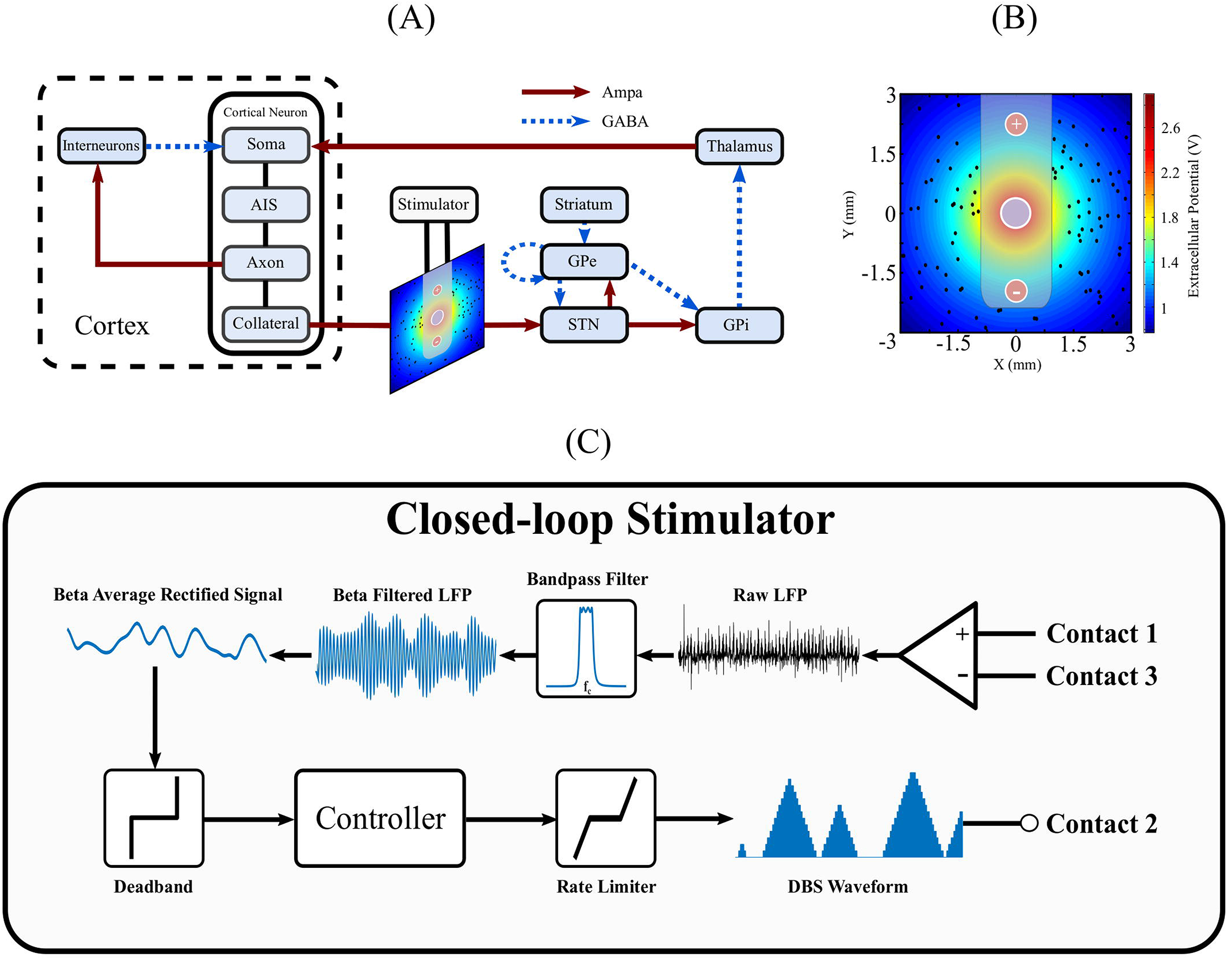
Schematic diagram of cortical basal ganglia network model. (A) Network diagram of cortical basal ganglia neuron populations. Excitatory and inhibitory synaptic connections within the network are represented as solid red arrow and blue dotted arrows, respectively. (B) Electric field distribution due to monopolar stimulation electrode. Cortical collaterals, represented as black dots, are oriented perpendicular to the page. The electrode bipolar recording contacts are represented by + and −, respectively. (C) Schematic diagram of the closed-loop stimulator utilizing an LFP derived measured of network beta-band activity. Contacts 1 and 3 represent the bipolar recording electrode contacts on the DBS electrode. The recorded LFP is bandpass filtered, rectified and averaged to calculate the average rectified value of the LFP beta-band activity. The beta average rectified value is used as input to a controller which determines an updated value for the DBS parameter being modulated. The updated DBS waveform is subsequently simulated at electrode contact 2 and varies the electric field distribution.

### 2.1 Network Structure

The major model components include single compartment, conductance-based biophysical models of cortical interneurons, STN, globus pallidus externa (GPe), globus pallidus interna (GPi), and thalamus neurons. Cortical Layer V pyramidal neurons, with axons projecting to the STN through the hyperdirect pathway, were simulated using multi-compartment, conductance-based biophysical models to enable extracellular application of DBS to cortical axon collaterals. The individual cell models have been validated and employed in previous modelling studies (Hahn and McIntyre, 2010; Kang and Lowery, 2013, 2014; Kumaravelu et al., 2016; Otsuka et al., 2004; Pospischil et al., 2008; Rubin and Terman, 2004; Terman et al., 2002). Six hundred cells consisting of one hundred STN, GPe, GPi, thalamic, cortical interneuron and cortical pyramidal neurons were connected through excitatory and inhibitory synapses, AMPA and GABAa, respectively, as described below, Figure 1 (a). While the type and direction of connections between the nuclei of the thalamo-cortico-basal ganglia network are well established, it is more difficult to ascertain the exact number of connections between individual neurons, and their relative strengths, in different nuclei. Input to a single neuron was therefore assumed to be from one or two neurons in each of the connected presynaptic nuclei, with the exception of the STN, which receives substantial direct cortical input (Nambu et al., 2002), with increased functional connectivity between the cortex and STN in the dopamine depleted state (West et al., 2018). Connections between neurons in the cortico-basal ganglia network followed a random connectivity pattern. Each STN neuron received excitatory input from five cortical neurons and inhibitory input from two GPe neurons. Each GPe neuron received inhibitory input from one striatal neuron, and one other GPe neuron, and excitatory input from two STN neurons. Each GPi neuron received excitatory input from a single STN neuron and inhibitory input from a single GPe neuron. Thalamic neuron received inhibitory input from a single GPi neuron. Cortical neurons received excitatory input from a single thalamic neurons and inhibitory input from ten interneurons. Interneurons received excitatory input from ten cortical neurons. The values for all model parameters are provided as Supplementary Material.

Pathological beta oscillations in the cortico-basal ganglia network were modelled based on the hypothesis that beta activity entering the network from the cortex is enhanced locally within the reciprocally coupled STN-GPe loop and propagates through the closed-loop network from the cortex through the basal ganglia, thalamus and back to cortex (Litvak et al., 2011; Mallet et al., 2008; Sharott et al., 2005). The strength of synaptic connections between the cortex and STN, between the STN and GPe, and the thalamus cortex were increased to induce beta oscillations within the network and the STN LFP, similar to that observed experimentally (West et al., 2018). This hypothesis is supported by studies investigating directional connectivity within the dopamine depleted cortico-basal ganglia, inversion of biophysical models using electrophysiological data from rats and patients (Marreiros et al., 2013; Moran et al., 2011; West et al., 2018), and functional imaging studies in individuals with Parkinson’s disease (Baudrexel et al., 2011; Fernández-Seara et al., 2015; Lalo et al., 2008). Previous simulation studies have similarly increased the strength of connections between nuclei to simulate the effects of dopamine depletion on basal ganglia network activity in Parkinson’s disease resulting in the emergence of oscillatory activity (Hahn and McIntyre, 2010; Humphries et al., 2006; Kang and Lowery, 2013; Kumaravelu et al., 2016; Nevado Holgado et al., 2010; Rubin and Terman, 2004).

### 2.2 Neuron Models

The compartmental membrane voltage of each neuron in the network is described by

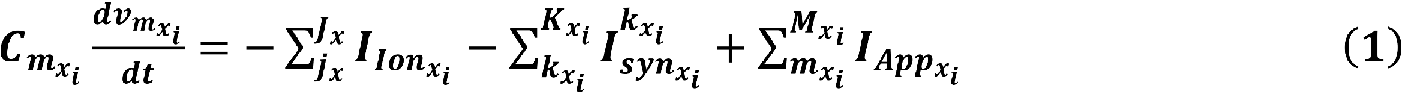

where *x* specifies the neuron population, *i* is the *i^th^* neuron in population *x*,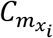 is the membrane capacitance of the *i^th^* neuron in population *x*,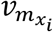 is the membrane potential of the *i^th^* neuron in population *x*. The membrane potential of the *i^th^* neuron in population *x* was calculated as the summation of the *J* ionic currents of population *x*’s neuron model, 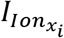, the *K_x_* synaptic currents which project to the*i^th^* neuron in population *x*, 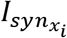, and the *M* intracellularly applied currents, 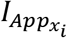 Further details regarding the neuron models are included below, and in the Supplementary Material.

#### 2.2.1 Cortex

The cortex was represented by a network of interneurons and cortical pyramidal neurons. The cortical neuron model, based on a layer V pyramidal tract neuron, comprised a soma, axon initial segment (AIS), main axon, and axon collateral as described by (Kang and Lowery, 2014). To summarize, the cortical neuron soma and interneuron models were based on the regular spiking neuron model developed in (Pospischil et al., 2008), while the model used to simulate the AIS, main axon, and axon collateral was based on results from the experimental modeling study in (Foust et al., 2011). The model compartments include leak, sodium, and three potassium ionic currents and an intracellular bias current for setting the neuron firing rate. The cortical soma compartment model excluded the D-type potassium current, while the AIS, main axon, and axon collateral compartments excluded the slow, voltage dependent potassium current. Cortical interneurons excluded both the D-type and slow, voltage dependent potassium currents.

#### 2.2.2 Subthalamic Nucleus

The STN model incorporated a physiological representation of STN neurons developed by (Otsuka et al., 2004) and implemented by (Hahn and McIntyre, 2010) that captures the generation of plateau potentials which have been identified as playing an important role in generating STN bursting activity that is observed during Parkinson’s disease (Beurrier et al., 1999). The STN model included leak, sodium, three potassium, two calcium ionic currents and an intracellular bias current for setting the neuron firing rate. Further details regarding the parameter values used can be found in the Supplementary Material and in (Hahn and McIntyre, 2010; Kang and Lowery, 2013, 2014; Otsuka et al., 2004).

#### 2.2.3 Globus Pallidus and Thalamus

GPe, GPi, and thalamic neurons were represented using the model developed in (Rubin and Terman, 2004) and implemented by (Hahn and McIntyre, 2010). The GPe and GPi neuron models included leak, sodium, two potassium, and two calcium ionic currents and an intracellular bias current for setting the neuron firing rates. GPe neurons included an additional intracellularly injected current to simulate the application of DBS to the GPe neuron model, assuming that an equivalent proportion of GPe neurons were stimulated to the proportion of extracellularly stimulated cortical neurons during DBS. Further details on the application of DBS to GPe neurons is included below in section 2.3. Thalamic neurons were modelled similarly, with the exception of excluding one of the calcium and one of the potassium currents. Striatal synaptic input to GPe neurons was modelled as a population of Poisson-distributed spike trains at 3 Hz.

#### 2.2.4 Synapses

Individual synaptic currents,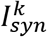, were described by

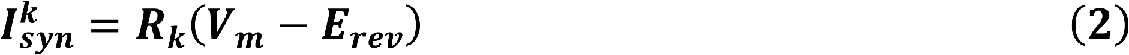

Where 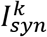 is the *k^th^* synaptic current, *R_k_* represents the kinetics of the onset decay of current following a presynaptic spike for synapse *k*, and *E_rev_* is the reversal potential for the appropriate type of synapse. Further details regarding the parameter values using in the synaptic models can be found in (Destexhe et al., 1994).

### 2.3 Application of DBS

The DBS electrode was modelled with three point source electrodes located in a homogeneous, isotropic medium of infinite extent and conductivity, *σ*, where a single point source was used to represent the application of extracellular DBS in a monopolar configuration, while the remaining two point source electrodes were used for simulating recording the local field potential with a bipolar, differential recording electrode. Propagation, inductive, and capacitive effects were assumed to be negligible, in accordance with the quasistatic approximation (Bossetti et al., 2008; Plonsey and Heppner, 1967).

The extracellular potential due DBS, 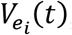, at each point located on the cortical collateral, *i*, located a distance *r*_i_ from the monopolar electrode was calculated as

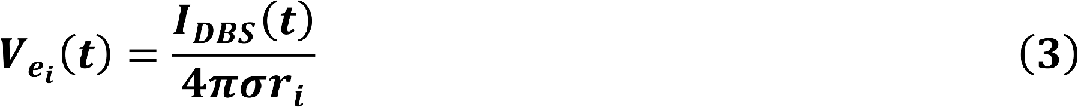

where σ is the conductivity of grey matter, with the specified value 0.27 S/m (Latikka et al., 2001), *I_DBS_* is the DBS current, simulated as a series of periodic cathodic rectangular current pulses of variable amplitude, frequency, and duration.

Cortical collaterals were randomly distributed around the monopolar electrode in a 6 mm by 6 mm square, using uniformly distributed random variables for their cartesian coordinates. The collaterals were oriented perpendicular to the cross-section, parallel to one another, and were not permitted to lie within the area covered by the cylindrical electrode lead of radius of 0.7 mm, Figure 1(B).

The application of DBS to the model was simulated by stimulating afferent STN projections resulting in antidromic activation of the cortex and GPe and orthodromic activation of excitatory and inhibitory afferent projections to the STN. This resulted in disruption of activity in the cortex and GPe and net inhibition of the STN, consistent with experimental observations (Filali et al., 2004; Li et al., 2012; Milosevic et al., 2018). The extracellular potential due to DBS was applied to cortical collaterals projecting to the STN from descending layer V pyramidal tract fibers (Kang and Lowery, 2014). It was assumed that an equal percentage of cortical and GPe neurons were activated during stimulation. During DBS, the percentage of activated cortical neurons was calculated and an intracellular DBS current was injected to the corresponding percentage of activated GPe neurons, where cortical neurons were labelled as activated during 130 Hz DBS if their collateral firing rate increased above 60 Hz. The entrainment order of the GPe neurons was generated as a randomized sequence from the first to the hundredth neuron in the population, where ten percent activation corresponded to the intracellular DBS current being delivered to the first ten GPe neurons in the entrainment order.

### 2.4 Local Field Potential Simulation

The STN LFP recorded at the bipolar, recording electrode was estimated as the summation of the extracellular potentials due to the spatially distributed synaptic currents across the STN population (Lindén et al., 2011). The *x* and *y* locations of STN neurons were randomly assigned as described previously, where the excitatory and inhibitory synapses for a given STN neuron were positioned at its *x* and *y* location, 250 μm from the bipolar electrode in the *z* direction. Assuming conduction within a purely resistive homogenous medium of infinite extent, the LFP at the bipolar electrode contacts was estimated as

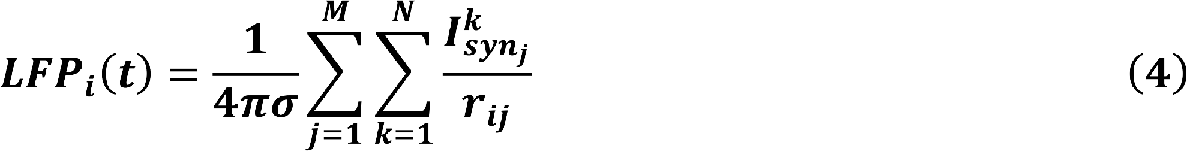

Where *LFP_i_(t)* is the LFP recorded by the *i^th^* bipolar, electrode contact at time *t*, 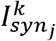 is the *k^th^* synaptic current of the *j^th^* STN neuron, and *r_ij_* is the distance from the *i^th^* electrode contact to the *k^th^* synapse of the *j^th^* STN neuron assuming M neurons, each with N synapses.

### 2.5 Beta-Band LFP Activity

The average rectified value (ARV) of the beta-band LFP was calculated by full-wave rectifying the filtered LFP signal using a fourth order Chebyshev band-pass filter with an 8 Hz bandwidth, centered about the peak in the LFP power spectrum. The last 100 ms epoch of the rectified signal was discarded to remove filtering artefact before taking the mean value of the last 100 ms epoch of the resulting signal. A target value for the beta ARV was estimated as the 20^th^ percentile of the beta ARV signal estimated for a thirty second epoch with DBS off. Cortical soma bias currents were modulated to vary the duration of beta activity within the network and simulated periods of high beta activity, or “beta bursts” periods, and low frequency activity. The duration of the beta bursts periods were varied to simulate short, “healthy bursts” of beta activity, *T_HB_,* and prolonged, “pathological bursts” of beta activity, *T_PB_*. Healthy burst periods were defined as 100 ms in duration while the duration of pathological bursts were drawn from a uniform distribution between 600 and 1000 ms to capture variability of pathological burst durations (Anidi et al., 2018; Tinkhauser et al., 2017a). The time between beta bursts, the interburst period, was fixed at 300 ms. The beta modulation signal was generated by selecting a random number at the start of each beta burst. If the random number was less than, or equal to 0.5 the burst was labeled healthy and its duration assigned as the healthy burst duration. If the random number was greater than 0.5 it was labeled pathological and its duration was set appropriately, selecting a value from the uniform distribution of pathological burst durations. During controller simulations, a beta ARV above the target corresponded to pathological beta activity, while a beta ARV below the target represented fluctuations of healthy beta activity. In practice, the target could be chosen based on an appropriate balance between symptom suppression and device power consumption. The controller input, *e*, at time *t* was calculated as the normalized error between the measured beta ARV, *b_measured_*, and the target beta ARV, *b_target_*, according to

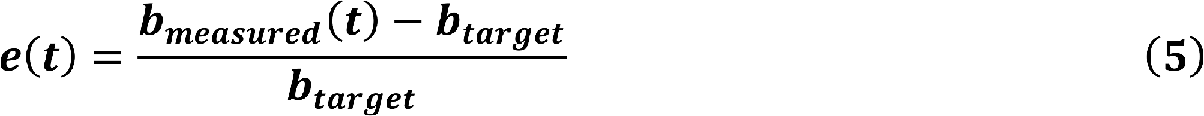

The controller operated with a sampling interval, *T_s_*, of 20 ms, updating the modulated DBS parameter at each controller call. During amplitude modulation, the DBS frequency and pulse duration were fixed at 130 Hz and 60 µs, respectively, with the amplitude varying between 0 – 3 mA, where the upper amplitude bound was selected as the amplitude which minimized the beta ARV. During frequency modulation, the DBS amplitude and pulse duration were fixed at 1.5 mA and 60 µs, respectively, with the frequency varying between 0 – 250 Hz.

It has been observed in clinical studies of closed-loop DBS amplitude modulation that rapid changes in the stimulation amplitude can potentially induce stimulation induced side-effects, or parathesias. To avoid unintentional parathesias Little *et al.* (2013) ramped their DBS amplitude from its minimum to maximum value over a 250 ms period during closed-loop DBS. Following this approach, the maximum tolerable rate limit for the modulated DBS parameter during closed-loop DBS was defined as

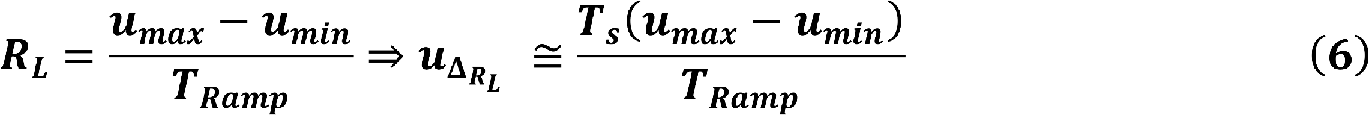

where *R_L_* is the rate limit of the DBS parameter per second, *T_Ramp_* is the duration of the ramping period, *T_s_* is the controller sampling period, 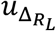 is the maximum tolerable variation of the DBS parameter per controller call, and *u_max_* and *u_min_* are the maximum and minimum bounds of the modulated DBS parameter. Utilizing this, the maximum rate limit for DBS amplitude modulation was calculated as R_L_ = 0.012 A/s and R_L_ = 1000 Hz/s for frequency modulation.

### 2.6 closed-loop Control

On-off, dual-threshold, P and PI controllers were investigated for closed-loop control of the DBS amplitude as detailed below. P and PI control were also used to investigate closed-loop control of the DBS frequency. The closed-loop DBS methodology simulated by the model is summarized in Figure 1(C).

#### 2.6.1 On-Off Controller

The on-off controller utilized a single target and increased or decreased the stimulation amplitude towards its upper or lower bounds if the beta ARV was measured above or below the target, respectively. The on-off controller is defined as

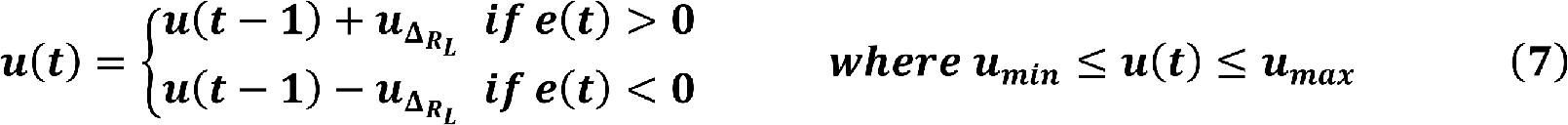

where *u(t)* is the modulated DBS parameter value, i.e. the stimulation amplitude, at time *t*,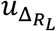 is the rate limit of the DBS parameter at each controller call, and *e(t)* is the controller error input signal at time *t*.

#### 2.6.2 Dual-Threshold Controller

The dual-threshold controller utilized a target range where the upper bound of the target range was selected as the 20^th^ percentile and the lower bound was selected as the 10^th^ percentile of the beta ARV with DBS off. If the beta ARV was greater than the upper bound of the target range, the error was calculated with respect to the upper bound, while if it was less than the lower target range bound, the error was calculated with respect to the lower bound. The behavior of the dual-threshold controller is defined as follows

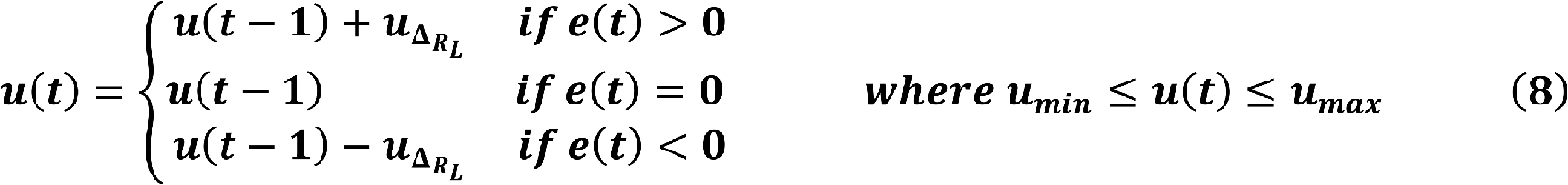

where the parameters are as described for the on-off controller.

#### 2.6.3 PI and P Controllers

The PI controller utilized a single target and is defined as

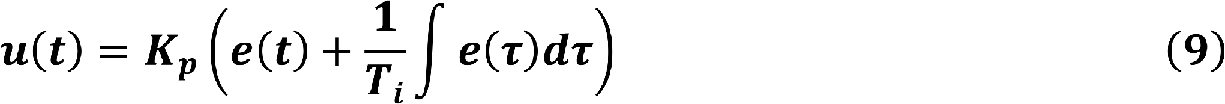

where *u(t)* is the modulated DBS parameter value at time *t*, *K_p_* is the controller proportional gain and *T_i_* is the controller integral time constant. The PI proportional component contributes to the modulated DBS parameter at time *t* by scaling the current controller measured error while the integral component contributes to the controller output at time *t* by scaling the integrated history of the controller measured errors up to time *t*. Conditional integration of the integral component was used to prevent integral wind-up, where integration of the integral component was paused if the modulated DBS parameter reached its upper or lower parameter bounds. Inclusion of a derivative gain, which would make the controller a PID controller rather than a PI controller, was deemed undesirable because below target fluctuations in beta-band activity, which occur during healthy beta bursts, would contribute to the modulated stimulation parameter through the derivative term. The P controller was simulated by omission of the integral term in (9).

#### 2.6.4 PI Controller Gain Tuning

The performance of PI controllers is heavily dependent on selection of appropriate values for the proportional gain, *K_p_*, and integral time constant, *T_i_*. The tuning process here is complicated by the constraint that the controller should not exceed the maximum tolerable rate limit of the modulated DBS parameter and that controller should act only on pathological beta bursts, while minimally effecting healthy beta bursts. The following tuning rules were thus designed for selecting PI controller parameters which adhere to these requirements.

##### 2.6.4.1 Selection of Integral Time Constant

The duration of beta bursts in the model varied between healthy and pathological durations, *T_HB_* and *T_PB_* respectively. It was thus desirable to select the integral time constant longer than the duration of healthy bursts and shorter than the duration of pathological beta bursts. Therefore, the integral time constant, *T_i_*, was selected as 0.2 s, so

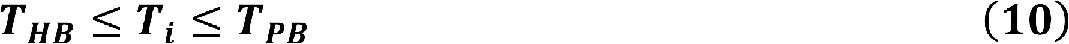

##### 2.6.4.2 Selection of Proportional Gain

The proportional gain, *K_p_*, was selected so that the rate limit of the modulated DBS parameter was not exceeded. This was calculated by differentiation of Eq. (9) and setting the DBS parameter rate limit, *R_L_*, as an inequality constraint

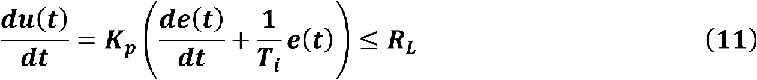

Rearranging, *K_p_* was defined as

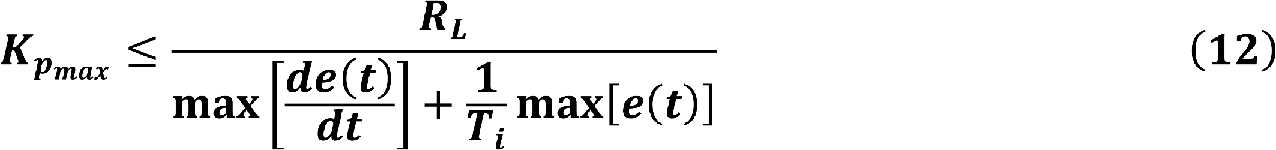

The maximum value of *e(t)* and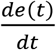 were estimated from a 30 second simulation with no DBS. Substituting in the corresponding values allows the calculation of an upper bound value for *K_p_*, 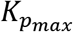. Using these rules, the PI controller parameters were calculated as (*K_p_, T_i_*) = (0.23, 0.20) for the PI amplitude controller and (*K_p_, T_i_*) = (19.30, 0.20) for the PI frequency controller.

A parameter sweep was conducted to select the proportional gain term for the P controllers, where the gain selected minimized the resulting mean controller error. The *K_p_* parameters were selected as 5.0 and 417 for the amplitude and frequency P controllers, respectively.

### 2.7 Simulation Details

The behavior of the model during continuous open-loop DBS was initially investigated with a constant level of beta activity within the network to characterize the relationship between the DBS waveform parameters and (i) the antidromic spike rate of cortical neurons detected at the cortical soma (ii) the beta-band power measured from the STN LFP and (iii) the firing rate of STN neurons. Ten simulations were conducted with initial random seeds varied between each simulation. Following this, a parameter sweep of the stimulation amplitude and frequency values was conducted to characterize the effect of parameter values on the LFP beta ARV. The parameter space was divided into 1024 linearly spaced sample points between the minimum and maximum bounds of the DBS amplitude, 0 – 3 mA, and frequency, 0 – 250 Hz. Each sample point corresponded to a 10 second simulation of open-loop DBS with the DBS parameters specified at that sample point. The performance of each closed-loop controller was then investigated in ten 30 second simulations where network beta activity was modulated as described in section 2.5. Ten independent beta modulation signals were generated, with each controller simulated for each modulation signal. The performance of the controllers were quantified in terms of the mean error of the half-wave rectified error signal and the mean power consumed, assuming a 1 KΩ electrode impedance, during each controller simulation. averaged over the ten controller simulations. A parameter sweep of the PI amplitude controller parameters was also conducted to investigate the effect of each parameter on the controller behavior. The sweep was conducted for *K_p_* values linearly spaced between (0, 6) and *Ti* values logarithmically spaced between (0, 6). All simulations were run from the model steady state, where an initial model simulation was run for 6 seconds to allow the network behavior to reach steady state, before the controller performance was then evaluated on the following 10 s. The initial model parameters in steady state were saved and used as the starting point for all subsequent simulations.

The model was simulated in the NEURON simulation environment (Hines and Carnevale, 1997) and implemented in Python using the PyNN API package (Davison, 2008). The model was integrated using a 0.01 ms timestep for all simulations. Simulations were run on the UCD Sonic high-performance computing cluster. Post-processing and signal analysis were done using custom scripts developed in MATLAB (The MathWorks, Inc., Natick, MA).

## 3 Results

The behavior of the model was first examined and compared with key features of the network behavior identified in experimental data from animal and human studies. Beta activity within the STN LFP, antidromic activation of cortical neurons and STN neural firing rates during continuous DBS with constant stimulation parameters were investigated. The firing rates of the cortical neurons were then modulated to simulate bursts of beta activity within the network and the performance of closed-loop DBS controllers to modulate either the DBS pulse amplitude or frequency were evaluated.

### 3.1 Network behavior during open-loop DBS

Cortical desynchronization and GPe entrainment were observed in the model during open-loop stimulation, Figure 2(D, F), with behavior qualitatively similar to DBS effects reported in experimental studies (Li et al., 2012; McConnell et al., 2012). Cortical antidromic firing rates matched well the observations of Li *et al*., (Li et al., 2012), where the rate of cortical antidromic spiking increased with increasing stimulation frequency to a maximum antidromic spike rate of 36.8±0.6 Hz at a stimulation frequency of approximately 130 Hz, Figure 2(A). Antidromic firing rate was defined as the number of successful stimulation-evoked antidromic activations detected at the soma of reliably stimulated cortical neurons where cortical collaterals were deemed to be ‘reliably’ stimulated when they were activated by at least 90 % of DBS pulses at the stimulation frequency. Further increasing the stimulation frequency resulted in a reduction in antidromic spike rate (Li et al., 2012), Figure 2(A). In the model, DBS influenced the cortical interneurons as the antidromic activation spread through the cortical network through branching collaterals. This altered activation of inhibitory interneurons through antidromic activation of the cortical neurons is consistent with the hypothesis proposed by Li *et al*., where this behavior was suggested as a potential mechanism for the failure of frequency following at higher frequencies (Li et al., 2012).

**Figure 2:**
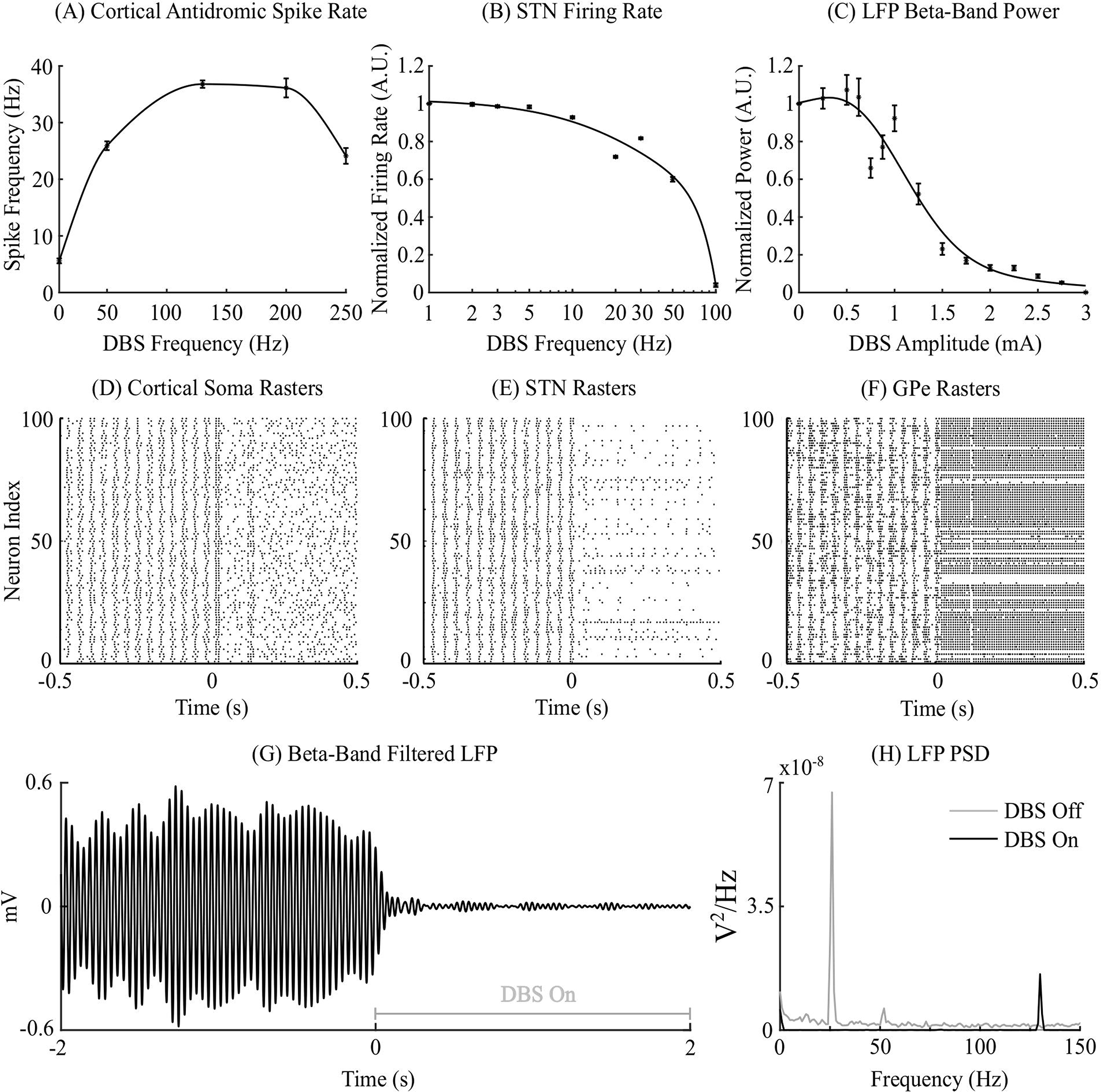
Cortico-basal ganglia network behavior during open-loop DBS. (A) Antidromic cortical spike rate during DBS with 2 mA amplitude, 60 μs pulse duration and varying frequency. (B) Normalized STN firing rate during DBS with 3 mA amplitude, 60 µs pulse duration, and varying DBS frequency. (C) Normalized STN LFP beta-band (22 – 30 Hz) power during DBS with 130 Hz frequency, 60 µs pulse duration, and varying DBS amplitude. (D-F) Cortical soma, STN and GPe population raster plots when DBS is off and during open-loop DBS with 2.5 mA amplitude, 130 Hz and 60 μs pulse. At time 0 s, DBS is applied to the network causing desynchronization of cortical somas, STN suppression and GPe entrainment (G) Simulated beta-band filtered LFP before and during stimulation with 2.5 mA amplitude, 130 Hz frequency and 60 μs pulse duration. DBS is turned off prior to time 0 s and switched on at 0 s. (H) STN LFP power spectral densities when DBS is off (grey line) and during DBS with 2.5 mA amplitude, 130 Hz frequency and 60 μs pulse duration (black line). When DBS is off the LFP has a peak at 26 Hz in the beta frequency band. When DBS is applied to the network the 26 Hz beta-band peak is suppressed and a peak appears in the LFP power spectrum at the stimulation frequency, 130 Hz.

Increasing the frequency of DBS resulted in a gradual reduction of the average STN neuron firing rate, Figure 2(B). Complete suppression of STN neurons was observed in the model at 100 Hz and is consistent with STN firing rate behavior reported by Milosevic *et al.* (2018) during DBS. In the model, differences in the properties of excitatory, AMPAergic, and inhibitory, GABAergic, synapses leads to a net inhibition of STN neurons at higher frequencies. In experimental studies, it has been suggested that inhibitory GABAergic afferents comprising the majority of terminals on the STN soma, in combination with differing rates of synaptic depletion, may explain observations of a reduction in STN firing rates during high frequency stimulation (Milosevic *et al.,* 2018).

The LFP beta-band power decreased non-linearly with increasing DBS amplitude, Figure 2(C). This relationship is similar to the reduction in LFP beta-band activity with increasing amplitude observed in clinical data which can be well-described by higher order models (Davidson et al., 2016). Low stimulation amplitudes had little influence on LFP beta-band activity with amplitudes less than 1.1 mA unable to suppress LFP beta-band power regardless of the stimulation frequency. The sensitivity of the amplitude of beta-band oscillations to the stimulation parameters is presented in Figure 3, where stimulation amplitudes above 1.1 mA reduced the LFP beta-band power for a broad range of stimulation frequencies above 40 Hz.

**Figure 3:**
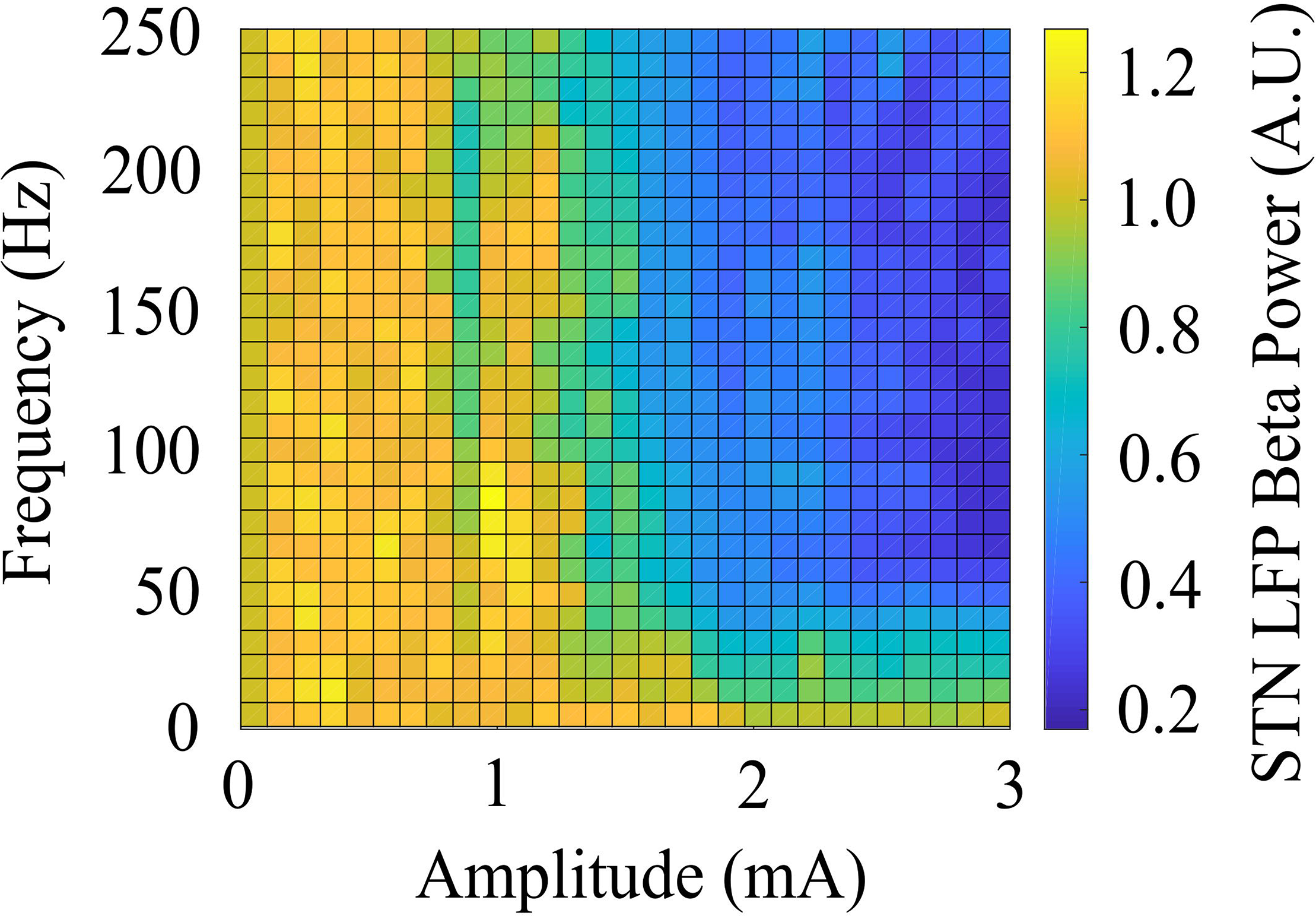
Effect of open-loop DBS parameters on STN LFP beta-band power. The STN LFP beta-band power was calculated during DBS with fixed 60 µs pulse duration and varying stimulation amplitude and frequency. The LFP beta-band power was normalized against the LFP beta-band power recorded when was DBS off. Stimulation amplitude and frequency values of 0 corresponded to the condition where DBS is off, resulting in an LFP beta-band power value of 1.

### 3.2 Closed-loop control of LFP beta-band activity

#### 3.2.1 Open-loop DBS

Model simulations with DBS off demonstrated modulation of the STN LFP beta-band activity, with varying periods of short and prolonged beta, Figure 4(A). The simulations without DBS were used to set a reference performance mean error value of 100 % for the controller. A reference value for the mean power consumption was obtained from simulations with open-loop, constant DBS at 2.5 mA, 130 Hz, and 60 µs pulse duration. This was used to set a reference mean power consumption value of 100 %, with a corresponding mean error value of approximately zero (0.4 %), Figure 4(B). These baseline performance values were used to compare the closed-loop DBS control strategies for keeping the beta ARV below the target ARV value.

**Figure 4:**
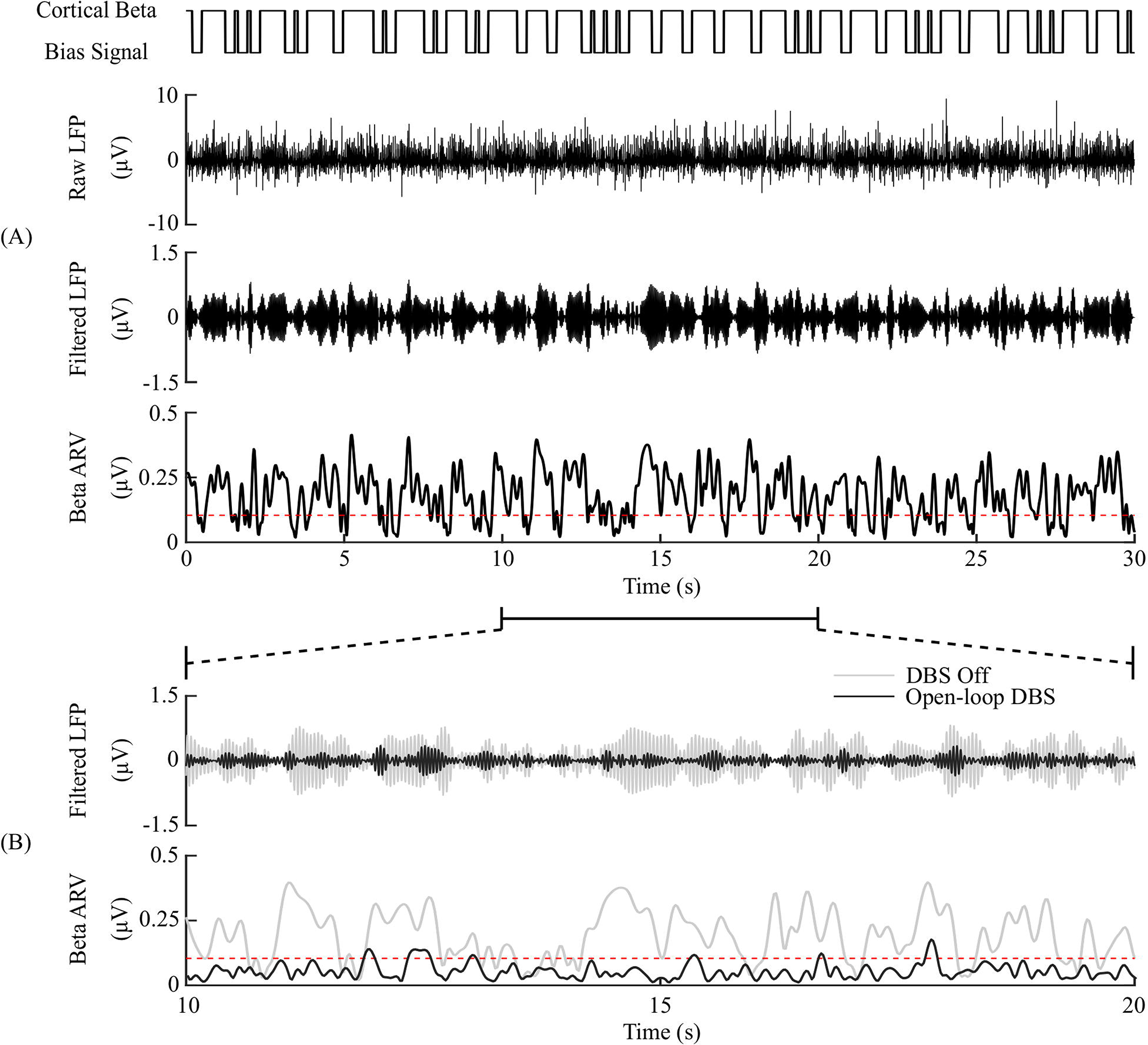
DBS off and open-loop DBS. (A) Example of a 30 s simulation with DBS off. The cortical bias current signal, top panel, represents the temporal modulation of the intracellular cortical bias current applied to the cortical neuron somas to generate activity in the beta frequency band. The raw LFP, beta-band filtered LFP and beta ARV are displayed in the next three panels. The target level for the beta ARV is represented by the red dotted line in the beta ARV figure panel. (B) 10 s simulation period of DBS off and open-loop DBS with 2.5 mA amplitude, 130 Hz frequency and 60 μs pulse duration. The panels correspond to the 10 – 20 s simulation period from (A). DBS off is represented as the grey lines in the filtered LFP and beta ARV panels, while open-loop DBS is displayed in black.

#### 3.2.2 Amplitude Modulation Controllers

The on-off controller resulted in a 63 % reduction in mean error compared to the DBS off condition, and a 60 % reduction in mean power consumed when compared with constant DBS, while the dual-threshold controller showed greater reduction in mean error with a 70 % decrease, but had a smaller mean power consumed reduction with a 50 % decrease, Figures 5 and 8.

**Figure 5:**
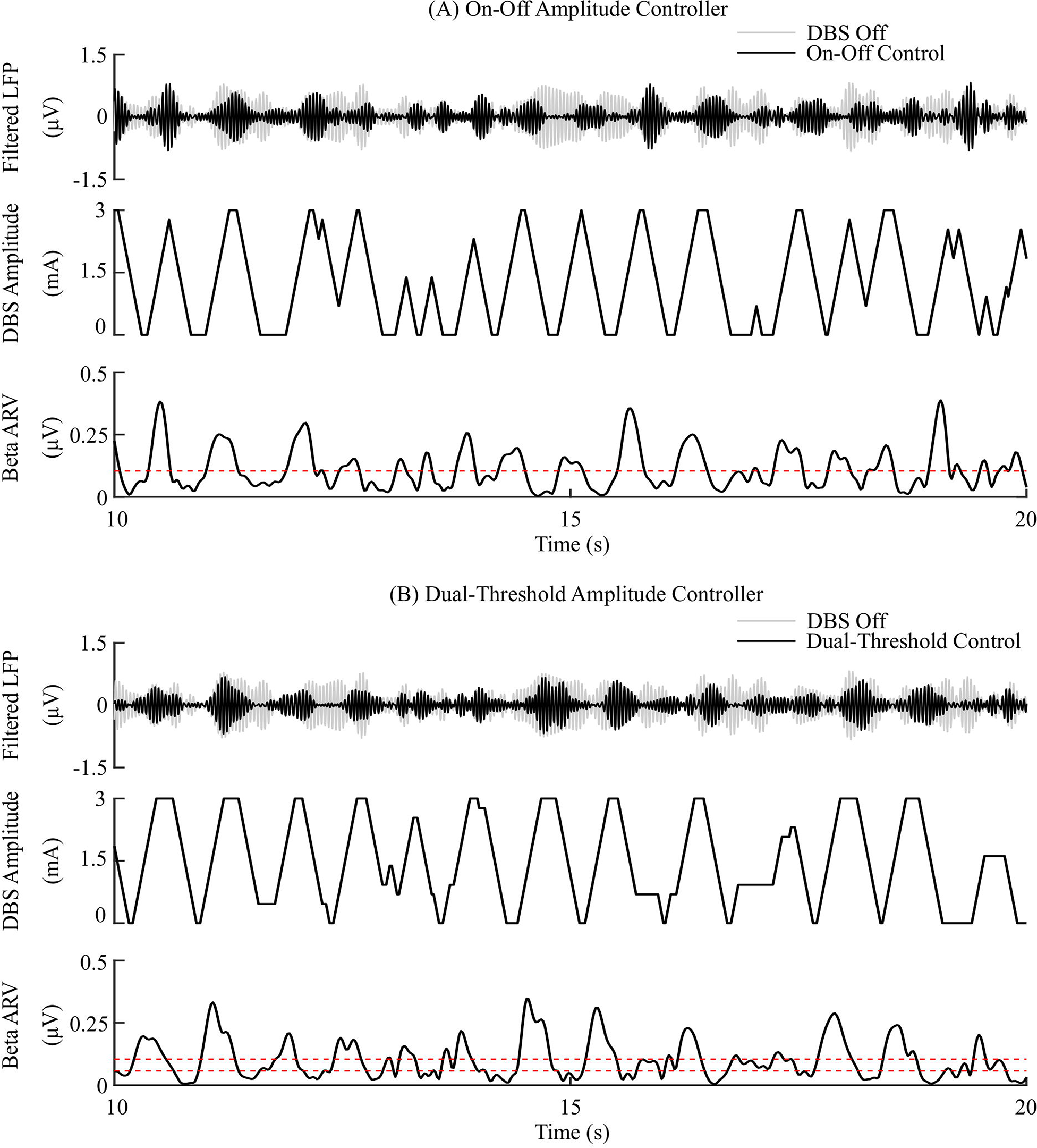
On-off and dual-threshold amplitude control, fixed 130 Hz frequency and 60 μs pulse duration. (A) On-off DBS amplitude controller. During simulation, the on-off controller increases or decreases the DBS amplitude by a fixed amount at each controller call, towards its upper or lower amplitude bounds, if the beta ARV is measured above or below the target value, the dotted red line in the beta ARV panel. (B) Dual-threshold amplitude controller. The dual-threshold controller uses a target beta ARV range, represented by the two dotted red lines. If the beta ARV is measured above the upper target range value or below the lower target range value the stimulation amplitude is increased or decreased, respectively, by a fixed amount towards the upper or lower bounds of the stimulation amplitude. If the beta ARV lies in the the target range the stimulation amplitude remains constant.

The P controller displayed a 62 % and 53 % reduction in the mean error and mean power consumed, while the PI controller showed a 79 % and 68 % decrease in the mean error and mean power consumed respectively, Figures 6 and 8.

**Figure 6:**
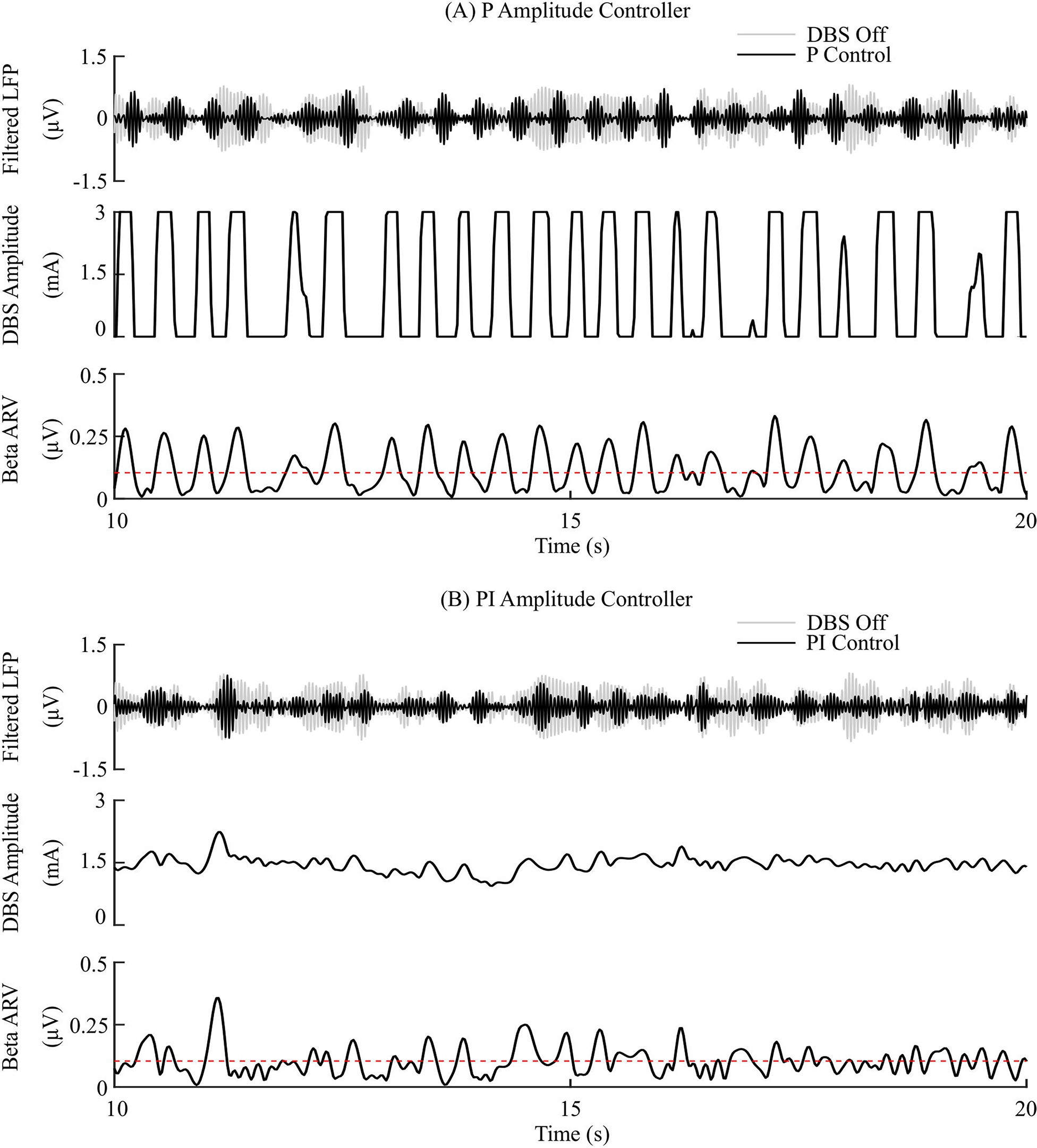
P and PI amplitude control, fixed 130 Hz frequency and 60 μs pulse duration. (A) P amplitude controller. At each controller call, the stimulation amplitude is calculated as the current measured error value scaled by the controller gain, *K_p_*. The controller stimulation amplitude is bounded between 0 – 3 mA, thus, negative error values correspond to turning DBS off. (B) PI amplitude controller. At each controller call, the stimulation amplitude is calculated as the summation of an integral term, i.e. the integration of the measured errors at previous controller calls, and the current measured error value, scaled by the integral time constant, *T_i_*, and the proportional gain, *K_p_*, respectively.

#### 3.2.3 Frequency Modulation Controllers

The P controller showed 72 % reduction in the mean error, with only a 1 % decrease in the mean power consumed when compared with continuous DBS. Better performance was obtained using the PI controller with a reduction in the mean error and mean power consumed by 83 % and 75 %, respectively, Figures 7 and 8.

**Figure 7:**
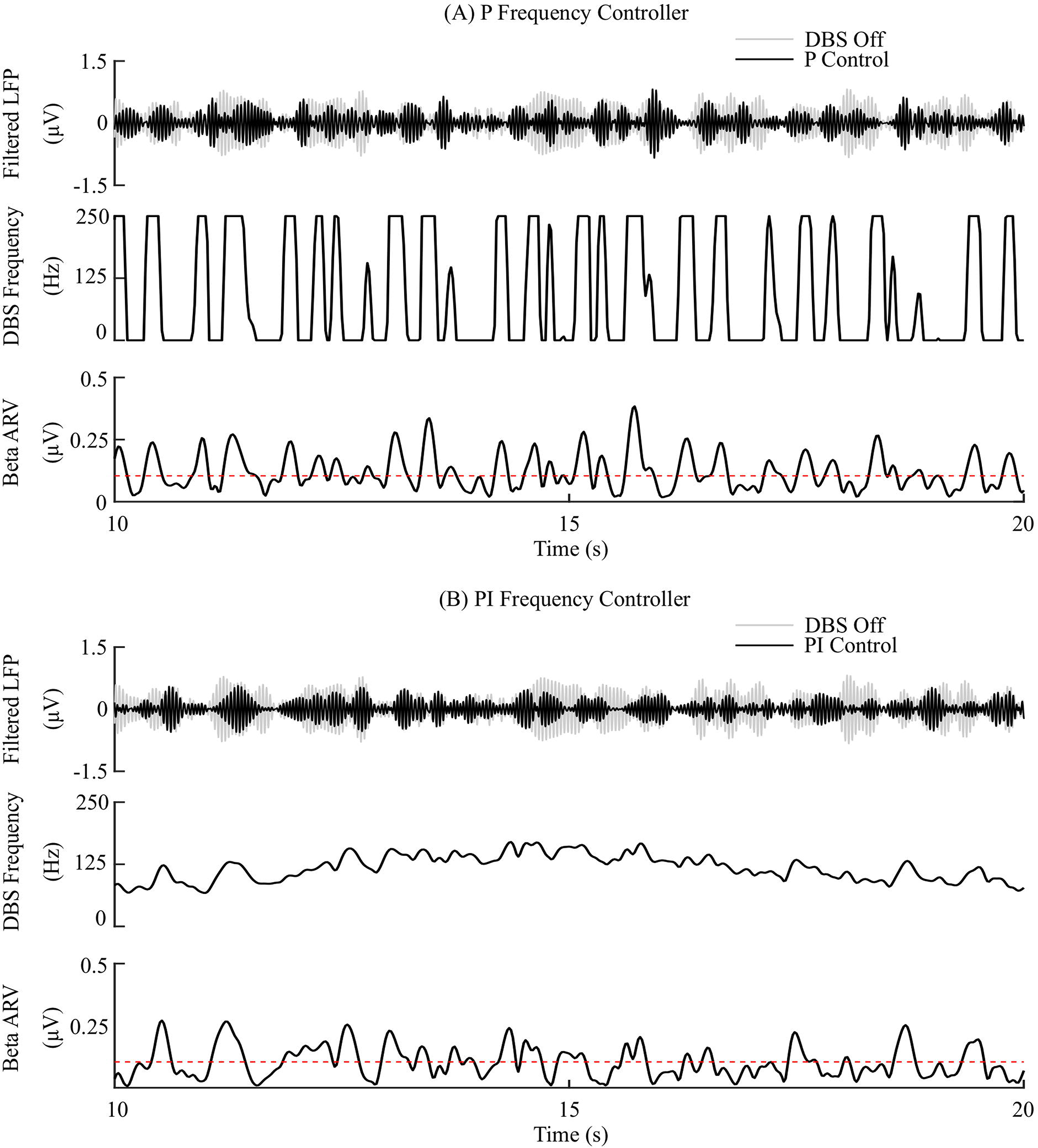
P and PI frequency control, fixed 1.5 mA amplitude and 60 μs pulse duration. (A) P frequency controller. At each controller call, the stimulation frequency is calculated as the current measured error value scaled by the controller gain, *K_p_*. The controller stimulation frequency is bounded between 0 – 250 Hz, thus, negative error values correspond to turning DBS off. (B) PI frequency controller. At each controller call, the stimulation frequency is calculated as the summation of an integral term, i.e. the integration of the measured errors at previous controller calls, and the current measured error value, scaled by the integral time constant, *T_i_*, and the proportional gain, *K_p_*, respectively.

**Figure 8:**
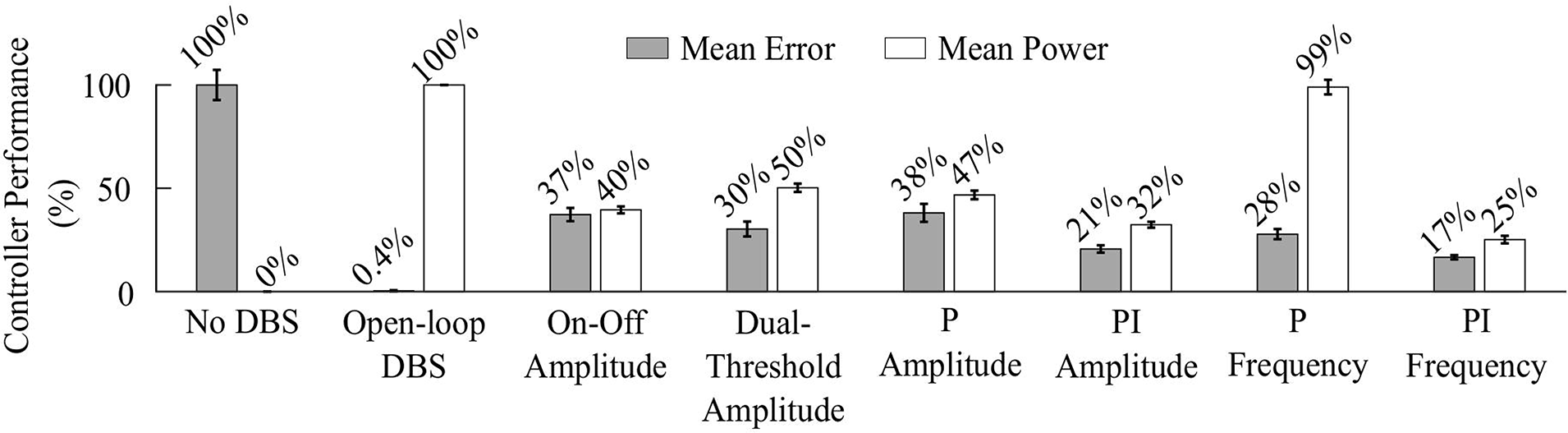
Summary of performance of closed-loop controllers. The normalized mean error (±std) and normalized mean power consumed (±std), averaged across ten 30 s simulations with ten independent beta modulation signals, of each closed-loop controller is presented. The mean error and mean power consumed are normalized against the mean error and mean power consumed when DBS is off or applied in open-loop with 2.5 mA amplitude, 130 Hz frequency and 60 μs pulse duration for each cortical beta modulation signal. The PI frequency controller performed best overall, reducing the mean error by 83% and the mean power consumed during stimulation by 75%.

The controller performances are summarized in Figure 8.

#### 3.2.4 Effect of varying PI parameter values

Having examined the PI controller using the derived parameters from the rule-tuning method, a sensitivity analysis was conducted to explore the parameters effect on controller performance. All the PI parameter value combinations tested as part of the controller parameter sensitivity analysis resulted in an approximately 55 % reduction in the mean error compared to DBS off. The mean power consumed showed a reduction of at least of 40 % for all combinations. A region of parameter space between *K_p_* = (0.25, 1) and *T_i_* = (0.02, 0.8) showed the greatest reduction in the mean error of 96 % and a 60 % reduction in power consumed at *K_p_* = 0.75 and *T_i_* = 0.19, Figure 9. A controller with relatively long *T_i_* and low *K_p_* resulted in slow performance, where the integral term slowly accumulated the error history and the modulated parameter varies slowly through the proportional term, Figure 9(C). In comparison, a controller with short *T_i_* and relatively large *K_p_* resulted in a fast controller response, where the error history accumulated quickly and the modulated parameter varied quickly between minimum and maximum values, Figure 9(E). PI parameter values selected using the tuning rule presented in this study resulted in a controller response which maintained the beta activity at the target level while adhering to rate constraints on the DBS amplitude, Figure 9(D).

**Figure 9:**
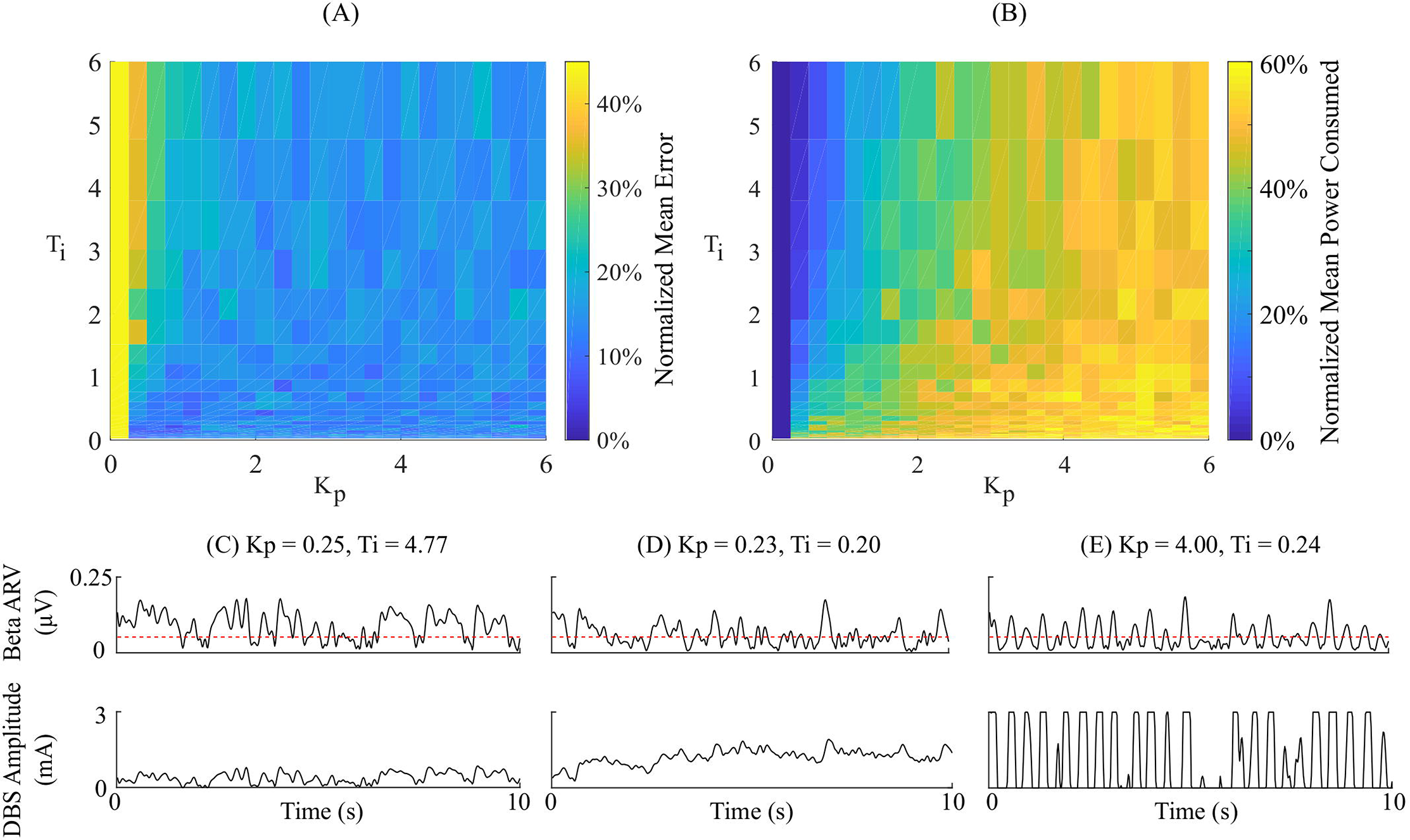
Effect of PI parameters on amplitude controller performance. (A) Normalized PI controller error vs. PI parameters. (B) Normalized PI controller power consumed vs. PI parameters. (C-E) Beta ARV and DBS amplitude due to varying *T_i_* and *K_p_* controller parameter values.

## 4 Discussion

A computational model is presented as an *in silico* testbed for developing and testing closed-loop DBS controllers designed to control LFP beta activity in PD. The model developed extends previous models by (i) incorporating the extracellular DBS electric field, (ii) captures both antidromic and orthodromic activation of afferent STN projections, (iii) simulates the synaptically generated STN LFPs and (iv) mimics temporal variation of network beta activity within the thalamo-cortico-basal ganglia loop. The model was first used to validate the performance of on-off and dual-threshold DBS amplitude closed-loop control schemes which have been tested clinically. P and PI controllers for modulating either the DBS amplitude or frequency were then investigated. PI controllers were found to outperform current clinically tested closed-loop controllers, displaying the greatest reductions in the controller mean error and power consumed during closed-loop DBS of the controllers examined.

### 4.1 Closed-loop Control of DBS

The observed reduction in power during on-off control within the model is consistent with clinical studies which have reported a 50 % reduction in mean power when compared with open-loop DBS (Little et al., 2013, 2016). In terms of the mean error, the dual-threshold controller performed better than on-off control, reducing the mean error by a further 7 %, Figure 8. This resulted in a 50 % reduction in the mean power consumed compared to open-loop DBS which is again in-line with the clinically reported 56.86 % reduction in energy delivered (Velisar et al., 2019). The improved performance maintaining the target beta level, at the cost of greater power consumption, was due to the dual-threshold controller’s ability to maintain a fixed stimulation amplitude when the beta ARV remained within its target bounds, Figure 5(B). Without this, the on-off controller results in a higher error but consumes less power during stimulation, Figures 5 and 8.

The mean error of P amplitude control was comparable to on-off control, while its mean power consumed was comparable to that of dual-threshold control, Figures 6(A) and 8. However, to achieve this performance, the P controller exceeded the prespecified rate limit of 0.012 A/s with a maximum rate observed of 0.150 A/s, exceeding clinically recommended limits to avoid side-effects. The P controller behaved similar to on-off control without rate limiting, or ‘bang-bang’ control, switching between its maximum and minimum values when the beta ARV was above or below the target. If a rate limiter is implemented on the P controller, it will behave similar to the on-off controller presented in this study, where deviations of the control variable from the target result in the amplitude varying by the maximum tolerable rate at each controller call. Arlotti *et al.* (2018) and Rosa *et al.* (2015) varied the stimulation voltage linearly, or proportionally, in response to the measured LFP beta-band power, rather than with respect to the error between the measured beta activity and a target as examined here. In that study, the control signal cannot have a value less than zero, while the control signal in this study is negative when beta activity is below the target. Due to the controller output bound at zero, DBS switches off when the control signal is negative here, while this behavior would not be observed when directly measuring the LFP beta-band power as the control signal. This distinction between using the beta activity or the error of the beta activity to a target is important to consider for clinical implementations of P controllers as this subtlety leads to disparate performances of the P controller.

The behavior of the P frequency controller was qualitatively similar to its amplitude counterpart, with the P frequency controller rapidly switching between its maximum and minimum values, Figures 7(A) and 8. Although the P frequency controller reduced the mean error by 72 %, the mean power consumed during stimulation was reduced by just 1 % when compared to open-loop DBS. The negligible change in mean power consumption was due to periods where the controller modulated the stimulation frequency between the minimum and maximum values of 0 Hz and 250 Hz. During these periods, the stimulation was either switched off or delivered close to double the number of stimulation pulses as during open-loop DBS or amplitude modulation, where the stimulation frequency was fixed at 130 Hz. With the stimulation amplitude fixed at 1.5 mA during frequency control this results in the same mean power consumed as open-loop DBS, Fig 8. The stimulation amplitude value selected for frequency control was chosen to allow use of the full span of stimulation frequencies, however it should be emphasized that by simply reducing this amplitude value or the controller’s upper frequency bound would result in the controller consuming less power.

The PI controllers for amplitude and frequency performed with 79 % and 83 % reductions in mean error, and a 68 % and 75 % decrease in mean power consumed for amplitude and frequency modulation, respectively, Figures 6(B), 7(B) and 8. The behaviors of both PI controllers were qualitatively similar, with the integral term increasing the modulated stimulation parameter to a value where it was effective at maintaining the beta ARV around the target level, Figures 6(B) and 7(B). Once at this value, fluctuations in the beta ARV resulted in proportional variation of the stimulation parameter to maintain the beta level. The integral term essentially overcomes the initial nonlinearity between the DBS parameter and the beta ARV, where a minimal value must be reached before the stimulation becomes effective. This is achieved by increasing the stimulation parameter to a region of parameter space where its relationship with the beta ARV is approximately linear, Figure 2(C) and Figure 3. The integral term varies the modulated stimulation parameter based on the error history in the system, whereas the on-off, dual-threshold and P controllers act only on the current error of the system at each controller call and thus have no memory of previous errors. For the on-off and dual-threshold controllers this can result in slow performance when the beta ARV exceeds the target and DBS is off. When this occurs, the DBS parameter must increase beyond the nonlinear region of its parameter space before stimulation becomes effective, which may take several controller calls. The gain of each P controller was selected as the gain value which minimized the mean error in a parameter sweep over the proportional gain values. The resulting P controllers were fast and essentially avoided the nonlinear region of the stimulation parameter space by quickly switching the stimulation parameter between its maximum and minimum values but did so at a rate that may be greater than is clinically desirable, Figures 6(A) and 7(A).

Overall, the PI frequency controller performed best, yielding the greatest reductions in mean error and mean power consumed of the controllers examined. Interestingly, the controller settled around a mean stimulation frequency of 125 Hz, which is in line with high frequency stimulation values utilized clinically. When modulating about this point the stimulation frequency varied between 80 – 160 Hz over the course of the simulation, with DBS remaining effective throughout the simulation, Figure 7(B). Clinical research has observed similar behavior where a 60 Hz DBS frequency was able to improve bradykinesia in PD patients (Blumenfeld et al., 2017). The authors hypothesized that 140 Hz high frequency stimulation and the lower frequency 60 Hz stimulation signals effectively decoupled the cortico-STN hyperdirect pathway during stimulation. The model presented in this study supports this hypothesis, with cortical desynchronization and STN firing rate suppression occurring during effective DBS, Figure 2(D, E). It is again important to note however, that due to the nonlinear relationship between DBS parameters and network beta activity there is a threshold stimulation amplitude value which must be reached before DBS frequency modulation becomes effective, in the model at approximately 1.1 mA, Figure 2(C).

A point of consideration for the controller results presented is that although the duration of LFP beta activity has been tested as a control variable for the on-off controller (Tinkhauser et al., 2017a), it has not been tested for either the dual-threshold or proportional controllers to date. Clinical studies investigating the dual-threshold and proportional controllers were limited to utilizing LFP beta band power as their control variables due to delays in the neurostimulator used during their studies (Arlotti et al., 2018; Velisar et al., 2019). This limitation is anticipated to be overcome in the next generation of neurostimulator devices and thus it will be feasible to utilize the duration of LFP beta activity as a control variable in the future (Velisar et al., 2019). With this in mind, the sampling frequency of controllers used in this study was selected so that fluctuations in the network beta band activity could be observed, with the controllers attempting to target only prolonged duration network beta activity.

#### 4.1.1 PI Controller Parameters

Suitable control parameters were identified using a rule-tuning approach which takes advantage of features of the biomarker that can be readily estimated clinically to derive suitable PI controller parameters, i.e. the threshold duration of pathological beta-band activity and constraints on the rate-of-change of stimulation parameters. When clinically tuning a PI controller for closed-loop DBS, the presented tuning rule could be used initially to coarsely-tune the controller, before further fine-tuning is achieved by varying the controller parameters using visual feedback of the modulated stimulation parameter. The intention here is to allow the clinician to further fine-tune the controller response if necessary, for example slowly increasing *K_p_* to increase the speed of the controller. Identifying suitable controller parameters could also be achieved in the model by utilizing an optimization technique and a suitable objective function, where the objective function captures the clinical considerations of the system. This approach, however, would require sampling multiple points in the parameter space which may not be practical clinically. An alternative controller design approach is to linearize the input-output relationship of the system using a model and subsequently design a controller which meets the required closed-loop system response (Liu et al., 2017; Santaniello et al., 2011; Su et al., 2019; Yang et al., 2018). This approach was used by Santaniello *et al.* (2011), Liu *et al.* (2017) and Su *et al.* (2019) where autoregressive models were derived from spiking neuron models. To normalize aberrant neural activity during parkinsonian tremor, Santaniello *et al.* (2011) designed a minimum variance controller, while Liu *et al.* (2017) implemented a generalized predictive control algorithm. In contrast, Su *et al.* (2019) optimized the parameters of a discrete PI controller to track a dynamic target of beta-band power which may be associated with fluctuations of the oscillatory activity during voluntary movement. Haddock *et al.* (2017) illustrated the potential of the approach by deriving an autoregressive model of the relationship between DBS amplitude and parkinsonian tremor from patient data, using the identified model as part of a model predictive controller for parkinsonian tremor (Haddock et al., 2017). The benefit of the autoregressive model approach is that derived models can be simulated in real-time and thus facilitate the use of advanced control techniques which require use of an internal model (Francis and Wonham, 1976). The disadvantage, however, is that it does not provide insight into the underlying physiological behavior of the system or its dynamics. Another drawback is that the identified model is valid only for the system operating region at which it was identified. Due to the dynamic, nonlinear nature of the parkinsonian neuromuscular system, identified models or controller parameters which were initially suitable during controller tuning may become unsuitable or provide suboptimal performance during different tasks, times throughout the day or as the disease progresses. Advanced adaptive techniques which automatically update autoregressive model coefficients or controller gains may be required to overcome this limitation (Cameron and Seborg, 1984; Chaillet et al., 2017; Santaniello et al., 2011). Nevertheless, the PI parameter rule-tuning approach presented in this study provides improved performance over currently tested closed-loop controllers, is simple to implement in a clinical setting and adheres to clinical considerations.

#### 4.1.2 Model Considerations

Previous modelling studies of closed-loop DBS have investigated LFP derived measures of network beta-band activity (Daneshzand et al., 2018; Popovych and Tass, 2019). Moreover, only a small number of models have simulated the coupled DBS electric field and network model during stimulation (Grant and Lowery, 2013; Santaniello et al., 2011). In the absence of a description of the electric field in the surrounding tissue during DBS, modelling studies are limited to simulating frequency modulation where the DBS waveform is injected as an intracellular current (Holt et al., 2016; Kang and Lowery, 2013; Popovych and Tass, 2019; Su et al., 2019). To develop clinically relevant closed-loop algorithms requires a model which captures modulation of the targeted networks behavior due to variations in both stimulation amplitude and frequency. The thalamocortical population model presented by Santaniello *et al.* (2011) incorporates both the DBS electric field and LFP simulation for investigating aberrant neural activity during parkinsonian tremor, but does not incorporate synaptic coupling between the neuron population, injecting a suprathreshold intracellular stimulus to neurons to drive spiking activity (Santaniello et al., 2011). As this modelling approach does not capture network interactions due to DBS it would, therefore, be unsuitable for modelling the full cortico-basal ganglia loop included here. Previous modelling studies of the cortico-basal ganglia during DBS have investigated network and DBS effects separately. Kang *et al.* (2013) and Kumaravelu *et al.* (2016) modelled the cortico-basal ganglia using networks of single compartment neuron models, however these models did not include simulation of extracellular DBS, the LFP or antidromic stimulation of afferent STN inputs. Grant *et al.* (2013) simulated extracellular DBS and the STN LFP, where the cortico-basal ganglia was modelled using a single compartment neuron model for the STN population and neural mass type models for the remaining network neuron populations. The model thus, does not capture complex network interactions such as antidromic activation of afferent STN inputs during stimulation. Antidromic activation of cortical afferent STN inputs and extracellular DBS was captured in a network model in (Kang and Lowery, 2014), however the model did not capture antidromic activation of GPe neurons during stimulation or simulation of the LFP. The model presented in this study builds on these previous modelling studies by incorporating extracellular DBS, STN LFP simulation, antidromic and orthodromic DBS effects and temporal variation of beta-band activity in a network model of the cortico-basal ganglia.

The controllers examined in this study represent the current landscape of clinically tested closed-loop DBS algorithms. The PI controller investigated is a natural extension of current state of the art closed-loop DBS research, and is the most commonly used control algorithm in industrial applications due to its robust performance in a wide range of operating conditions and its functional simplicity. PID-type controllers have been investigated in previous modelling studies of closed-loop DBS for PD (Gorzelic et al., 2013; Su et al., 2019). However, these studies utilized control variables which are not readily accessible during clinical studies, where Gorzelic *et al.* (2013) investigated using both thalamic reliability and GPi synaptic conductance as control variables and Su *et al.* (2019) used the beta-band power of GPi neuron spike times. Thus, direct comparisons between these studies and clinical research is difficult. The model presented here utilizes an LFP derived measure of network beta-band oscillatory activity analogous to that employed during clinical closed-loop DBS research, and thus facilitates a direct comparison between the performance of controllers tested in the model and in clinical research.

The purpose of the model is to provide an *in silico* testbed for developing and testing closed-loop DBS strategies which can be directly related to clinical closed-loop DBS research. Although this study focused on using an LFP derived measure of network beta-band activity for closed-loop DBS there is extensive research in identifying alternative biomarkers for PD symptoms and stimulation side-effects, such as entropy (Anderson et al., 2015; Dorval et al., 2010; Dorval and Grill, 2014; Fleming and Lowery, 2019; Syrkin-Nikolau et al., 2017), phase-amplitude coupling (de Hemptinne et al., 2013; De Hemptinne et al., 2015), coherence (Al-Fatly, 2019) and gamma-band activity (Swann et al., 2016, 2018) based measures. A restriction of the presented model is that it does not capture the neural mechanisms which lead to parkinsonian tremor, a hallmark symptom of PD, and is thus unsuited for investigating tremor-based closed-loop DBS (Helmich, 2018; Hirschmann et al., 2017). However, with this in mind, it is anticipated that future controllers which employ alternative methods or advanced techniques, such as low frequency stimulation (Blumenfeld et al., 2017; Fasano and Lozano, 2014) or phase-based (Holt et al., 2016, 2019; Tass, 2003; Tass et al., 2012) and linear-delayed feedback methods (Popovych and Tass, 2019), may still be applicable when alternative biomarkers are implemented as control variables.

### 4.2 Limitations

While the model captures several key features of the parkinsonian cortico-basal ganglia during DBS, it remains an approximation of the true system. Due to the limited access to the cortico-basal ganglia structures and data available in literature, the individual neuron models are based on parameters recorded from animal models of the disease from separate studies. Kumaravelu *et al.* (2016) used experimental data from 6-OHDA lesioned rats to parameterize their network model. In the presented model, synaptic coupling was tuned, and the cortical population was biased to induce increased beta-band oscillatory activity within the network, with striatal input to the basal ganglia network being simplified as a population of poisson-distributed spike trains. Although research suggests that the striatum has an influence on beta-band oscillatory activity within the network (Feingold et al., 2015; McCarthy et al., 2011) investigation of its effects on network oscillatory activity was not the purpose of this paper, and thus its contribution to the network was simplified. A recent study investigating the role of exogenous cortical and striatal beta inputs to the STN-GPe network using detailed multi-compartment models of STN and GPe showed that resonant beta-band oscillatory activity within the STN-GPe loop becomes phase-locked to exogenous cortical beta inputs and that this behavior can be further promoted by striatal input to the loop with the correct phase (Koelman and Lowery, 2019). The network presented here captures the exogenous cortical patterning of the STN-GPe loop but omits possible further amplification of the beta-band oscillatory activity due to the striatum. A consequence of the simplification of the cortical and cortico-striatal networks, and the inability to accurately capture all of the complex network interactions which lead to elevated beta-band activity in PD is that the oscillatory activity reemerges relatively quickly post-stimulation. In clinical studies it is observed that STN beta-band power shows long-lasting attenuation post-stimulation, with attenuation dependent on the stimulation duration (Bronte-Stewart et al., 2009; Temperli et al., 2003). This behavior is not captured by the model, where beta-band activity reemerges quickly when DBS is off or ineffective. Finally, the electrode was simulated as a point source electrode within an ideal homogeneous resistive volume conducting medium of infinite extent. Computational studies have previously demonstrated the point source approximation to be a valid prediction of the activation of a population of neurons during DBS when calculating the number and spatial distribution of neurons activated around the electrode (Zhang and Grill, 2010). However, in reality, the electrode geometry, its encapsulation tissue, and the capacitive and dispersive electrical properties of the tissue can have a substantial effect on the electric field distribution and on the properties of the simulated LFP (Lempka and McIntyre, 2013) and the activation thresholds of target neurons during DBS (Grant and Lowery, 2010; McIntyre et al., 2004). More realistic geometrical and anatomical properties of the tissues could be incorporated through coupling of the model to anatomically-realistic finite element models. In keeping with studies regarding the spatial reach of the LFP, it was assumed that the LFP signal was dominated by synaptic currents from neurons in the vicinity of the recording electrode (Lempka and McIntyre, 2013; Lindén et al., 2011), however, a contribution from more distal neurons outside of the STN network is also possible. The spatial distribution of synapses within the dendritic structures and neuron morphology can further influence the LFP, however these were not considered here.

## 5 Conclusion

A computational model of closed-loop control of DBS for PD is presented that simulates (i) the extracellular DBS electric field, (ii) antidromic and orthodromic activation of STN afferent fibers, (iii) the LFP detected at non-stimulating contacts on the DBS electrode and (iv) temporal variation of beta-band activity within the cortico-basal ganglia network. The model captures experimentally reported network behavior during open-loop DBS and provides an *in silico* testbed for developing novel, clinically-relevant closed-loop control strategies for updating either the amplitude or frequency of DBS. Clinically tested on-off and dual-threshold amplitude controllers were examined and exhibited reductions in power consumption comparable with their clinically reported performance. A new rule-tuning method for selecting PI controller parameters to target prolonged, pathological duration beta-band oscillatory activity whilst adhering to clinical constraints was developed. The resulting performance of both amplitude and frequency PI controllers outperformed the current clinically investigated on-off and dual-threshold closed-loop amplitude control strategies in terms of both power consumption and their ability to maintain the LFP derived measure of network beta-band activity at a target value. As the available technology progresses towards a new generation of closed-loop or adaptive stimulators, it is likely that testing novel control algorithms in computational models, such as those presented here, will become a valuable first step prior to clinical testing in patients.

## Supporting information

Supplemental Model Details

6

## 7 Author Contributions

All experiments were performed in the Neuromuscular Systems Laboratory in University College Dublin, Ireland. JF and ML: designed study experiments, interpreted results of experiments, prepared the figures, edited and revised the manuscript, and approved the final version of the manuscript. JF: performed experiments and analyzed data and drafted the manuscript. ED and ML: conceived preliminary experiment design.

## 10 Supplementary Material

Full details of cortico-basal ganglia network model equations are included in supplementary material.

## 11 Data Availability Statement

The datasets generated for this study are available on request to the corresponding author, while the model source code will be made available from ModelDB (https://senselab.med.yale.edu/modeldb/) upon manuscript publication.

