## Supplemental Model Details for "Simulation of closed-loop deep brain stimulation control schemes for suppression of pathological beta oscillations in Parkinson’s disease"

### *Supplementary Material*

#### 1 Supplementary Data

The transmembrane potential of each neuron ($v_m$) from each population in the network model is described by a Hodgkin-Huxley type neuron model. Details of each neuron model are described below.

##### 1.1 STN Neuron Model

The STN model used is from (Otsuka et al., 2004) using the implementation from Hahn and McIntyre (2011). The membrane potential of an STN neuron is described by

$$C_m \frac{dv_m}{dt} = -I_l - I_{Na} - I_K - I_A - I_L - I_T - I_{Ca-K} - \sum_k I_{syn}^k + I_{app} \qquad (A.1)$$

where $C_m$ is the membrane capacitance, $I_l$ is the leak current, $I_{Na}$ is a sodium current, $I_K$ is a Kv3-type potassium current, $I_A$ is a voltage dependent A-type potassium current, $I_L$ is an L-type long lasting calcium current, $I_{Ca\text{-}K}$ is a calcium activated potassium current, $I_{syn}$ are synaptic currents and $I_{app}$ is the neuron bias current. The equations governing the ionic currents are detailed in Table 1.

**Table 1: STN Neuron Model Equations**

| Current | Equation |
|---|---|
| $\boldsymbol{I_l}$ | $g_l(v_m - E_l)$ |
| $\boldsymbol{I_{Na}}$ | $g_{Na}m^3h(v_m - E_{Na})$ |
| $\boldsymbol{I_K}$ | $g_K n^4(v_m - E_K)$ |
| $\boldsymbol{I_A}$ | $g_A a^2 b(v_m - E_K)$ |
| $\boldsymbol{I_L}$ | $g_L c^2 d_1 d_2(v_m - E_{Ca})$ |
| $\boldsymbol{I_T}$ | $g_T p^2 q(v_m - E_{Ca})$ |
| $\boldsymbol{I_{Ca-K}}$ | $g_{Ca-K} r^2(v_m - E_K)$ |

The gating kinetics of the ionic conductances are governed by

$$\frac{dx}{dt} = \frac{x_\infty - x}{\tau_x} \qquad (A.2)$$

where $x$ stands for $a$, $b$, $c$, $d_1$, $d_2$, $h$, $m$, $n$, $p$, $q$, or $r$. The steady state activation and inactivation functions are given by

$$x_\infty = \frac{1}{1 + e^{\frac{v+\theta_x}{k_x}}} \qquad (A.3)$$

where $\theta_x$ and $k_x$ are the half-activation/half-inactivation voltage and slopes, respectively. The activation time constants are given by

$$\tau_x = \tau_x^0 + \frac{\tau_x^1}{e^{-\frac{(v+\theta_x^1)}{\sigma_x^1}} + e^{-\frac{(v+\theta_x^2)}{\sigma_x^2}}} \qquad (A.4)$$

The dynamics of the intrinsic calcium level $[Ca]_i$ are described by

$$\frac{d[Ca]_i}{dt} = \epsilon_{Ca}\left(-\frac{I_T + I_L}{2F} - k_{Ca}[Ca]_i\right) \qquad (A.5)$$

where $[Ca]_i$ is the intrinsic calcium concentration, $\epsilon_{Ca}$ is a constant which accounts for the combined effects of intracellular calcium buffering mechanisms and cellular geometry, $F$ is Faraday's constant and $k_{Ca}$ is the calcium pump rate. Parameter values for the STN neuron model are summarized in Table 2.

**Table 2: STN Neuron Model Parameters**

| Parameter | Values | Parameter | Values | Parameter | Values |
|---|---|---|---|---|---|
| $\boldsymbol{C_m}$ | 1 μF/cm$^2$ | $\theta_r$ | 0.17e-3 mM | $\tau_r$ | 2 ms |
| $\boldsymbol{L}$ | 60 μm | $k_a$ | -14.7 mV | $\theta_a^\tau$ | -40 mV |
| $\boldsymbol{Diam}$ | 60 μm | $k_b$ | 7.5 mV | $\theta_b^{\tau 1}, \theta_b^{\tau 2}$ | -60, -40 mV |
| $\boldsymbol{g_l}$ | 0.35 mS/cm$^2$ | $k_c$ | -5 mV | $\theta_c^{\tau 1}, \theta_c^{\tau 2}$ | -27, 50 mV |
| $\boldsymbol{g_{Na}}$ | 49 mS/cm$^2$ | $k_{d_1}$ | 7.5 mV | $\theta_{d_1}^{\tau 1}, \theta_{d_1}^{\tau 2}$ | -40, -20 mV |
| $\boldsymbol{g_K}$ | 57 mS/cm$^2$ | $k_{d_2}$ | 0.02 μM | $\theta_h^{\tau 1}, \theta_h^{\tau 2}$ | -50, -50 mV |
| $\boldsymbol{g_A}$ | 5 mS/cm$^2$ | $k_h$ | 6.4 mV | $\theta_m^\tau$ | -53 mV |
| $\boldsymbol{g_L}$ | 15 mS/cm$^2$ | $k_m$ | -8 mV | $\theta_n^{\tau 1}, \theta_n^{\tau 2}$ | -40, -40 mV |
| $\boldsymbol{g_T}$ | 5 mS/cm$^2$ | $k_n$ | -14 mV | $\theta_p^{\tau 1}, \theta_p^{\tau 2}$ | -27, -102 mV |
| $\boldsymbol{g_{Ca-K}}$ | 1 mS/cm$^2$ | $k_p$ | -6.7 mV | $\theta_q^{\tau 1}, \theta_q^{\tau 2}$ | -50, -50 mV |
| $\boldsymbol{E_l}$ | -60 mV | $k_q$ | 5.8 mV | $\sigma_a$ | -0.5 mV |
| $\boldsymbol{\theta_a}$ | -45 mV | $k_r$ | -0.08 μM | $\sigma_b^1, \sigma_b^2$ | -30, 10 mV |
| $\boldsymbol{\theta_b}$ | -90 mV | $\tau_a^0, \tau_a^1$ | 1, 1 ms | $\sigma_c^1, \sigma_c^2$ | -20, 15 mV |
| $\boldsymbol{\theta_c}$ | -30.6 mV | $\tau_b^0, \tau_b^1$ | 0, 200 ms | $\sigma_{d_1}^1, \sigma_{d_1}^2$ | -15, 20 mV |
| $\boldsymbol{\theta_{d_1}}$ | -60 mV | $\tau_c^0, \tau_c^1$ | 45, 10 ms | $\sigma_h^1, \sigma_h^2$ | -15, 16 mV |
| $\boldsymbol{\theta_{d_2}}$ | 0.1 μM | $\tau_{d_1}^0, \tau_{d_1}^1$ | 400, 500 ms | $\sigma_m$ | -0.7 mV |
| $\boldsymbol{\theta_h}$ | -45.5 mV | $\tau_h^0, \tau_h^1$ | 0, 24.5 ms | $\sigma_n^1, \sigma_n^2$ | -40, 50 mV |

| | | | | | |
|---|---|---|---|---|---|
| $\boldsymbol{\theta_m}$ | -40 mV | $\tau_m^0, \tau_m^1$ | 0.2, 3 ms | $\sigma_p^1, \sigma_p^2$ | -10, 15 mV |
| $\boldsymbol{\theta_n}$ | -41 mV | $\tau_n^0, \tau_n^1$ | 0, 11 ms | $\sigma_q^1, \sigma_q^2$ | -15, 16 mV |
| $\boldsymbol{\theta_p}$ | -56 mV | $\tau_p^0, \tau_p^1$ | 5, 0.33 ms | $I_{app}$ | -0.125 nA |
| $\boldsymbol{\theta_q}$ | -85 mV | $\tau_q^0, \tau_q^1$ | 0, 400 ms | | |

where *L* and *Diam* represent the length and diameter of the single compartment STN neuron.

#### 1.2 Globus Pallidus Neuron Model

The GPe and GPi models used are from (Rubin and Terman, 2004; Terman et al., 2002) using the implementation from Hahn and McIntyre (2011). The membrane potential of an GP neuron is described by

$$C_m \frac{dv_m}{dt} = -I_l - I_{Na} - I_K - I_T - I_{Ca} - I_{AHP} - \sum_k I_{syn}^k + I_{app} \qquad (A.6)$$

where $C_m$ is the membrane capacitance, $I_l$ is the leak current, $I_{Na}$ is a sodium current, $I_K$ is a potassium current, $I_T$ is a low-threshold T-type calcium current, $I_{Ca}$ is a high-threshold calcium current, $I_{AHP}$ is a calcium activated, voltage-independent 'afterhyperpolarization' potassium current, $I_{syn}$ are synaptic currents and $I_{app}$ is the neuron bias current. The equations governing the ionic currents are detailed in Table 1.

**Table 3: GP Neuron Model Equations**

| Current | Equation |
|---|---|
| $\boldsymbol{I_l}$ | $g_l(v_m - E_l)$ |
| $\boldsymbol{I_{Na}}$ | $g_{Na}m^3h(v_m - E_{Na})$ |
| $\boldsymbol{I_K}$ | $g_K n^4(v_m - E_K)$ |
| $\boldsymbol{I_T}$ | $g_T p^2 q(v_m - E_{Ca})$ |
| $\boldsymbol{I_{AHP}}$ | $g_{AHP} r^2(v_m - E_K)$ |

The gating kinetics of the ionic conductances are governed by

$$\frac{dx}{dt} = \frac{x_\infty - x}{\tau_x}$$

where *x* stands for *h, m, n, p, q,* or *r*. The steady state activation and inactivation functions are given by

$$x_\infty = \frac{1}{1 + e^{(v+\theta_x)/k_x}}$$

where $\theta_x$ and $k_x$ are the half-activation/half-inactivation voltage and slopes, respectively. The activation time constants are given by

$$\tau_x = \tau_x^0 + \frac{\tau_x^1}{e^{-\frac{(v+\theta_x^1)}{\sigma_x^1}} + e^{-\frac{(v+\theta_x^2)}{\sigma_x^2}}} \quad (A.7)$$

Parameter values for the GP neuron model are summarized in Table 4:

**Table 4: GP Neuron Model Parameters**

| Parameter | Values | Parameter | Values | Parameter | Values |
|---|---|---|---|---|---|
| $\boldsymbol{C_m}$ | 1 µF/cm$^2$ | $\theta_q$ | -85 mV | $\tau_r$ | 2 ms |
| $\boldsymbol{L}$ | 60 µm | $\theta_r$ | 0.17 µM | $\theta_h^{\tau 1}, \theta_h^{\tau 2}$ | -50, -50 mV |
| $\boldsymbol{Diam}$ | 60 µm | $k_h$ | 6.4 mV | $\theta_m^{\tau}$ | -53 mV |
| $\boldsymbol{g_l}$ | 0.35 mS/cm$^2$ | $k_m$ | -7 mV | $\theta_n^{\tau 1}, \theta_n^{\tau 2}$ | -40, -40 mV |
| $\boldsymbol{g_{Na}}$ | 49 mS/cm$^2$ | $k_n$ | -14 mV | $\theta_p^{\tau 1}, \theta_p^{\tau 2}$ | -27, -102 mV |
| $\boldsymbol{g_K}$ | 57 mS/cm$^2$ | $k_p$ | -6.7 mV | $\theta_q^{\tau 1}, \theta_q^{\tau 2}$ | -50, -50 mV |
| $\boldsymbol{g_T}$ | 5 mS/cm$^2$ | $k_q$ | 5.8 mV | $\sigma_h^1, \sigma_h^2$ | -15, 16 mV |
| $\boldsymbol{g_{AHP}}$ | 1 mS/cm$^2$ | $k_r$ | -0.08 µM | $\sigma_m$ | -0.7 mV |
| $\boldsymbol{E_l}$ | -60 mV | $\tau_h^0, \tau_h^1$ | 0, 4.5 ms | $\sigma_n^1, \sigma_n^2$ | -40, 50 mV |
| $\boldsymbol{\theta_h}$ | -45.5 mV | $\tau_m^0, \tau_m^1$ | 0.001, 0.1 ms | $\sigma_p^1, \sigma_p^2$ | -10, 15 mV |
| $\boldsymbol{\theta_m}$ | -38 mV | $\tau_n^0, \tau_n^1$ | 0, 2.4 ms | $\sigma_q^1, \sigma_q^2$ | -15, 16 mV |
| $\boldsymbol{\theta_n}$ | -42 mV | $\tau_p^0, \tau_p^1$ | 5, 0.33 ms | $I_{app_{GPe}}$ | -0.009 nA |
| $\boldsymbol{\theta_p}$ | -56 mV | $\tau_q^0, \tau_q^1$ | 0, 400 ms | $I_{app_{GPi}}$ | 0.006 nA |

where *L* and *Diam* represent the length and diameter of the single compartment GP neurons. The calcium dynamics for the GP neurons are the model the same as for the STN neuron in equation A.5. GPe and GPi neurons are modelled using the same parameter values in Table 4 with the exception of their bias currents, $I_{app_{GPe}}$ and $I_{app_{GPi}}$.

#### 1.3 Thalamic Neuron Model

The thalamic neuron model is from (Rubin and Terman, 2004) where the membrane potential of a thalamic neuron is described by

$$C_m \frac{dv_m}{dt} = -I_l - I_{Na} - I_K - I_T - \sum_k I_{syn}^k \quad (A.8)$$

where $C_m$ is the membrane capacitance, $I_l$ is the leak current, $I_{Na}$ is a sodium current, $I_K$ is a potassium current, $I_T$ is a low-threshold T-type calcium current and $I_{syn}$ are synaptic currents. The equations governing the ionic currents are detailed in Table 5.

**Table 5: Thalamic Neuron Model Equations**

| Current | Equation |
|---|---|
| $\boldsymbol{I_l}$ | $g_l(v_m - E_l)$ |
| $\boldsymbol{I_{Na}}$ | $g_{Na}m_\infty^3 h(v_m - E_{Na})$ |
| $\boldsymbol{I_K}$ | $g_K\big(0.75 * (1-h)\big)^4(v_m - E_K)$ |
| $\boldsymbol{I_T}$ | $g_T p_\infty^2 q(v_m - E_{Ca})$ |

The gating kinetics of the ionic conductances are governed by

$$\frac{dx}{dt} = \frac{x_\infty - x}{\tau_x} \qquad (A.9)$$

where $x$ stands for $h$ or $q$. The steady state activation and inactivation functions are given by

$$m_\infty = \frac{1}{1 + e^{-(v-37)/7}} \qquad (A.10)$$

$$h_\infty = \frac{1}{1 + e^{(v+41)/4}} \qquad (A.11)$$

$$p_\infty = \frac{1}{1 + e^{-(v+60)/6.2}} \qquad (A.12)$$

$$r_\infty = \frac{1}{1 + e^{(v+84)/4}} \qquad (A.13)$$

and the activation time constants are

$$\tau_h = \frac{1}{a_h + b_h} \qquad (A.14)$$

$$\tau_r = 28 + e^{-(v+25)/10.5} \qquad (A.15)$$

with specific functions $a_h$ and $b_h$

$$a_h = 0.128e^{-(v+46)/18} \qquad (A.16)$$

$$b_h = \frac{4}{1 + e^{-(v+23)/5}} \qquad (A.17)$$

Parameter values for the thalamic neuron model are summarized in Table 6:

**Table 6: Thalamic Neuron Model Parameters**

| Parameter | Values |
|---|---|
| $\boldsymbol{C_m}$ | 100 μF/cm$^2$ |
| $\boldsymbol{L}$ | 100 μm |
| $\boldsymbol{Diam}$ | 100 μm |
| $\boldsymbol{g_l}$ | 5 mS/cm$^2$ |
| $\boldsymbol{g_{Na}}$ | 300 mS/cm$^2$ |
| $\boldsymbol{g_K}$ | 500 mS/cm$^2$ |
| $\boldsymbol{g_T}$ | 500 mS/cm$^2$ |
| $\boldsymbol{E_l}$ | -60 mV |
| $\boldsymbol{E_{Na}}$ | 50 mV |
| $\boldsymbol{E_K}$ | -90 mV |
| $\boldsymbol{E_{Ca}}$ | 0 mV |

where *L* and *Diam* represent the length and diameter of the single compartment thalamic neuron.

#### 1.4 Cortical Pyramidal Neuron Model

The cortical pyramidal neuron model is modelled as a multicompartment neuron with the cortical soma compartment based on a model presented in (Pospischil et al., 2008) and the cortical axon compartments based on the model presented in (Foust et al., 2011). The membrane potential of each compartment in the cortical neuron model is described by the general equation

$$C_{m_x}\frac{dv_{m_x}}{dt} = -I_{l_x} - I_{Na_x} - I_{K_x} - I_{Kd_x} - I_{M_x} - \sum_k I_{syn_x}^k + I_{app_x} \qquad (A.18)$$

where *x* is the neuron compartment identifier, $C_m$ is the membrane capacitance, $I_l$ is the leak current, $I_{Na}$ is a sodium current, $I_K$ is a potassium current, $I_{Kd}$ is a D-type potassium current, $I_M$ is a slow, voltage-dependent potassium current, $I_{syn}$ are synaptic currents and $I_{app}$ is the neuron bias current. The equations governing the ionic currents are detailed in Tables 7 and 9.

##### 1.4.1 Cortical Pyramidal Neuron Soma Model

The equations governing the ionic currents are detailed in Table 7 below.

**Table 7: Cortical Pyramidal Neuron Soma Model Equations**

| Current | Equation |
|---|---|
| $\boldsymbol{I_{l_{soma}}}$ | $g_{l_{soma}}\left(v_{m_{soma}} - E_{l_{soma}}\right)$ |
| $\boldsymbol{I_{Na_{soma}}}$ | $g_{Na_{soma}} m^3 h\left(v_{m_{soma}} - E_{Na_{soma}}\right)$ |

| | |
|---|---|
| $\boldsymbol{I_{K_{soma}}}$ | $g_{K_{soma}} n^4 \left(v_{m_{soma}} - E_{K_{soma}}\right)$ |
| $\boldsymbol{I_{M_{soma}}}$ | $g_M p \left(v_m - E_{K_{soma}}\right)$ |

The gating kinetics of ionic conductances *m*, *h* and *n* are derived from a first order kinetic scheme and governed by

$$\frac{dx}{dt} = \alpha_x(1 - x) - \beta_x x \tag{A.19}$$

where x stands for either *m*, *h* or *n*. The steady state activation functions and the time constants for *m*, *h* and *n* are given by

$$x_\infty = \frac{\alpha_x}{\alpha_x + \beta_x} \tag{A.20}$$

$$\tau_x = \frac{1}{\alpha_x + \beta_x} \tag{A.21}$$

where

$$\alpha_m = \frac{-0.32\left(v_{m_{soma}} - v_T - 13\right)}{e^{-\frac{v_{m_{soma}} - v_T - 13}{4}} - 1} \tag{A.22}$$

$$\beta_m = \frac{0.28\left(v_{m_{soma}} - v_T - 40\right)}{e^{-\frac{v_{m_{soma}} - v_T - 40}{5}} - 1} \tag{A.23}$$

$$\alpha_h = 0.128 e^{-\frac{v_{m_{soma}} - v_T - 17}{18}} \tag{A.24}$$

$$\beta_h = \frac{4}{1 + e^{-\frac{v_{m_{soma}} - v_T - 40}{5}}} \tag{A.25}$$

$$\alpha_n = \frac{-0.32\left(v_{m_{soma}} - v_T - 15\right)}{e^{-\frac{v_{m_{soma}} - v_T - 15}{5}} - 1} \tag{A.26}$$

$$\beta_n = 0.5 e^{-\frac{v_{m_{soma}} - v_T - 10}{40}} \tag{A.27}$$

The gating kinetics of the slow non-inactivating potassium current $I_M$ is governed by

$$\frac{dp}{dt} = \frac{p_\infty - p}{\tau_p} \tag{A.28}$$

and the steady state activation function and time constant is given by

$$p_{\infty} = \frac{1}{1 + e^{-\frac{(v_{m_{soma}}+35)}{10}}} \qquad (A.29)$$

$$\tau_p = \frac{\tau_{max}}{3.3e^{\frac{v_{m_{soma}}+35}{20}} + e^{-\frac{v_{m_{soma}}+35}{20}}} \qquad (A.30)$$

Parameter values for the cortical pyramidal neuron soma are summarized in Table 8:

**Table 8: Cortical Pyramidal Neuron Soma Model Parameters**

| Parameter | Values |
|---|---|
| $\boldsymbol{C_{m_{soma}}}$ | 1 µF/cm$^2$ |
| $\boldsymbol{L_{soma}}$ | 35 µm |
| $\boldsymbol{Diam_{soma}}$ | 25 µm |
| $\boldsymbol{g_{l_{soma}}}$ | 0.1 mS/cm$^2$ |
| $\boldsymbol{g_{Na_{soma}}}$ | 50 mS/cm$^2$ |
| $\boldsymbol{g_{K_{soma}}}$ | 5 mS/cm$^2$ |
| $\boldsymbol{g_{M_{soma}}}$ | 0.07 mS/cm$^2$ |
| $\boldsymbol{E_l}$ | -70 mV |
| $\boldsymbol{E_{Na}}$ | 50 mV |
| $\boldsymbol{E_K}$ | -100 mV |
| $\boldsymbol{v_T}$ | -55 mV |
| $\boldsymbol{\tau_{max}}$ | 1000 ms |
| $\boldsymbol{I_{app_{soma}}}$ | 0.245 nA |

where $L_{soma}$ and $Diam_{soma}$ represent the length and diameter of the soma compartment and the soma bias current, $\boldsymbol{I_{app_{soma}}}$, value sets the cortical neuron firing rate within the beta-frequency band.

##### 1.4.2 Cortical Pyramidal Neuron Axon Model

The equations governing the ionic currents for the cortical axon initial segment (AIS), axon myelinated segments, nodes of Ranvier segments and collateral segments are detailed in Table 9 below.

**Table 9: Cortical Pyramidal Neuron Axon Model Equations**

| Current | Equation |
|---|---|
| $\boldsymbol{I_{l_{seg_{id}}}}$ | $g_{l_{seg_{id}}}\left(v_{m_{seg_{id}}} - E_l\right)$ |
| $\boldsymbol{I_{Na_{seg_{id}}}}$ | $g_{Na_{seg_{id}}} m^3 h \left(v_{m_{seg_{id}}} - E_{Na}\right)$ |

| | |
|---|---|
| $\boldsymbol{I_{K_{seg_{id}}}}$ | $g_{K_{seg_{id}}} n \left( v_{m_{seg_{id}}} - E_K \right)$ |
| $\boldsymbol{I_{Kd_{seg_{id}}}}$ | $g_{Kd_{seg_{id}}} pq \left( v_{m_{seg_{id}}} - E_K \right)$ |

where $seg_{id}$ specifies the axon compartment type. The gating kinetics of ionic conductances *m, h, n, p and q* are governed by

$$\frac{dx}{dt} = \frac{x_\infty - x}{\tau_x} \qquad (A.31)$$

where the steady state activation functions and time constants are described by

$$\tau_m = \frac{1}{\alpha_m + \beta_m}, \qquad m_\infty = \frac{\alpha_m}{\alpha_m + \beta_m} \qquad (A.32)$$

$$\tau_h = \frac{1}{\alpha_h + \beta_h}, \qquad h_\infty = \frac{1}{1 + e^{(v_{m_{seg_{id}}} + 60)/6.2}} \qquad (A.33)$$

$$\tau_n = \frac{1}{\alpha_n + \beta_n}, \qquad n_\infty = \frac{\alpha_n}{\alpha_n + \beta_n} \qquad (A.34)$$

$$\tau_p = 1\, ms, \qquad p_\infty = 1 - \frac{1}{1 + e^{(v_{m_{seg_{id}}} - V^p_{1/2})/q_p}} \qquad (A.35)$$

$$\tau_r = 1500\, ms, \qquad r_\infty = 1 - \frac{1}{1 + e^{(v_{m_{seg_{id}}} - V^r_{1/2})/q_r}} \qquad (A.36)$$

$$\alpha_m = \phi \frac{0.182 \left( v_{m_{seg_{id}}} + 30 \right)}{1 - e^{-(v_{m_{seg_{id}}} + 30)/8}}, \qquad \beta_m = -\phi \frac{0.124 \left( v_{m_{seg_{id}}} + 30 \right)}{1 - e^{(v_{m_{seg_{id}}} + 30)/8}} \qquad (A.37)$$

$$\alpha_h = \phi \frac{0.028 \left( v_{m_{seg_{id}}} + 45 \right)}{1 - e^{-(v_{m_{seg_{id}}} + 45)/6}}, \qquad \beta_h = -\phi \frac{0.0091 \left( v_{m_{seg_{id}}} + 70 \right)}{1 - e^{(v_{m_{seg_{id}}} + 70)/6}} \qquad (A.38)$$

$$\alpha_n = \phi \frac{0.01 \left( v_{m_{seg_{id}}} - 30 \right)}{1 - e^{-(v_{m_{seg_{id}}} - 30)/9}}, \qquad \beta_n = -\phi \frac{0.002 \left( v_{m_{seg_{id}}} - 30 \right)}{1 - e^{(v_{m_{seg_{id}}} - 30)/9}} \qquad (A.39)$$

$$\phi = Q_{10}^{T - 23/10} \qquad (A.40)$$

where the $Q_{10}$ effect, captured by $\phi$, regulates temperature dependence, with $Q_{10} = 2.3$. Parameter values for the cortical axon model compartments are summarized in Table 10.

**Table 10: Cortical Pyramidal Neuron Axon Model Parameters**

| Parameter | Values | Parameter | Values | Parameter | Values |
|---|---|---|---|---|---|
| $C_{m_{AIS}}$ | 0.8 µF/cm$^2$ | $g_{l_{AIS}}$ | 0.033 mS/cm$^2$ | $g_{Kd_{AIS}}$ | 15 mS/cm$^2$ |
| $C_{m_{Myelin}}$ | 0.04 µF/cm$^2$ | $g_{l_{Myelin}}$ | 0 mS/cm$^2$ | $g_{Kd_{Myelin}}$ | 0 mS/cm$^2$ |
| $C_{m_{Node}}$ | 0.8 µF/cm$^2$ | $g_{l_{Node}}$ | 20 mS/cm$^2$ | $g_{Kd_{Node}}$ | 7.2 mS/cm$^2$ |
| $C_{m_{Collateral}}$ | 0.8 µF/cm$^2$ | $g_{l_{Collateral}}$ | 0.033 mS/cm$^2$ | $g_{Kd_{Collateral}}$ | 0.6 mS/cm$^2$ |
| $L_{AIS}$ | 20 µm | $g_{Na_{AIS}}$ | 400 mS/cm$^2$ | $E_l$ | -70 mV |
| $L_{Myelin}$ | 500 µm | $g_{Na_{Myelin}}$ | 1 mS/cm$^2$ | $E_{Na}$ | 60 mV |
| $L_{Node}$ | 2 µm | $g_{Na_{Node}}$ | 280 mS/cm$^2$ | $E_K$ | -90 mV |
| $L_{Collateral}$ | 500 µm | $g_{Na_{Collateral}}$ | 133.33 mS/cm$^2$ | $V_{1/2}^{p}$ | -43 mV |
| $Diam_{AIS}$ | 1.2 µm | $g_{K_{AIS}}$ | 2 mS/cm$^2$ | $V_{1/2}^{r}$ | -67 mV |
| $Diam_{Myelin}$ | 1.4 µm | $g_{K_{Myelin}}$ | 0 mS/cm$^2$ | $q_p$ | 8 |
| $Diam_{Node}$ | 1.2 µm | $g_{K_{Node}}$ | 0.5 mS/cm$^2$ | $q_r$ | 7.3 |
| $Diam_{Collateral}$ | 0.5 µm | $g_{K_{Collateral}}$ | 1 mS/cm$^2$ | | |

#### 1.5 Cortical Interneuron Model

The cortical interneuron model used was the single-compartment model from (Pospischil et al., 2008). The membrane potential of the cortical interneuron model is described by

$$C_m \frac{dv_m}{dt} = -I_l - I_{Na} - I_K - \sum_k I_{syn}^k + I_{app} \qquad (A.41)$$

where $C_m$ is the membrane capacitance, $I_l$ is the leak current, $I_{Na}$ is a sodium current, $I_K$ is a potassium current, $I_{syn}$ are synaptic currents and $I_{app}$ is the neuron bias current. The equations governing the ionic currents are detailed below in Tables 11.

**Table 11: Cortical Interneuron Model Equations**

| Current | Equation |
|---|---|
| $I_l$ | $g_l(v_m - E_l)$ |
| $I_{Na}$ | $g_{Na}m^3h(v_m - E_{Na})$ |
| $I_K$ | $g_K n^4(v_m - E_K)$ |

The gating kinetics of ionic conductances *m*, *h* and *n* are derived from a first order kinetic scheme and governed by

$$\frac{dx}{dt} = \alpha_x(1-x) - \beta_x x \qquad (A.42)$$

where x stands for either *m*, *h* or *n*. The steady state activation functions and the time constants for *m*, *h* and *n* are given by

$$x_\infty = \frac{\alpha_x}{\alpha_x + \beta_x} \qquad (A.43)$$

$$\tau_x = \frac{1}{\alpha_x + \beta_x} \qquad (A.44)$$

where

$$\alpha_m = \frac{-0.32(v_m - v_T - 13)}{e^{-\frac{v_m - v_T - 13}{4}} - 1} \qquad (A.45)$$

$$\beta_m = \frac{0.28(v_m - v_T - 40)}{e^{-\frac{v_m - v_T - 40}{5}} - 1} \qquad (A.46)$$

$$\alpha_h = 0.128 e^{-\frac{v_m - v_T - 17}{18}} \qquad (A.47)$$

$$\beta_h = \frac{4}{1 + e^{-\frac{v_m - v_T - 40}{5}}} \qquad (A.48)$$

$$\alpha_n = \frac{-0.32(v_m - v_T - 15)}{e^{-\frac{v_m - v_T - 15}{5}} - 1} \qquad (A.49)$$

$$\beta_n = 0.5 e^{-\frac{v_m - v_T - 10}{40}} \qquad (A.50)$$

Parameter values for the cortical interneurons are summarized in Table 12:

**Table 12: Cortical Interneuron Model Parameters**

| Parameter | Values |
|---|---|
| $\boldsymbol{C_m}$ | 1 μF/cm$^2$ |
| $\boldsymbol{L}$ | 35 μm |
| $\boldsymbol{Diam}$ | 25 μm |
| $\boldsymbol{g_l}$ | 0.15 mS/cm$^2$ |
| $\boldsymbol{g_{Na}}$ | 50 mS/cm$^2$ |
| $\boldsymbol{g_K}$ | 10 mS/cm$^2$ |

| | |
|---|---|
| $\boldsymbol{E_l}$ | -70 mV |
| $\boldsymbol{E_{Na}}$ | 50 mV |
| $\boldsymbol{E_K}$ | -100 mV |
| $\boldsymbol{v_T}$ | -55 mV |
| $\boldsymbol{I_{app_{soma}}}$ | 0.07 nA |

where *L* and *Diam* represent the length and diameter of the cortical interneuron.

#### 1.6 Network Synaptic Connectivity

The details of the synaptic connections in the network are detailed in Table 13, where the number of connections between each presynaptic and postsynaptic population, the strength of the synaptic connections and the transmission delay are specified:

**Table 13: Network Synaptic Connectivity Parameters**

| Pre | Post | # Connections | $w_{syn}$ (µS) | $t_{syn}$ (ms) |
|---|---|---|---|---|
| Cortical - Axon Node | Cortical - Interneuron | 10 | (0, 2.5e-3) | 2 |
| Cortical - Interneuron | Cortical - Soma | 10 | (0, 6.0e-3) | 2 |
| Cortical - Axon Collateral | STN Neuron | 5 | 0.12 | 1 |
| STN Neuron | GPe Neuron | 1 | 0.11 | 4 |
| GPe Neuron | GPe Neuron | 1 | 0.015 | 4 |
| GPe Neuron | STN Neuron | 2 | 0.11 | 3 |
| Striatal Neuron | GPe Neuron | 1 | 0.01 | 1 |
| STN Neuron | GPi Neuron | 1 | 0.11 | 2 |
| GPe Neuron | GPi Neuron | 1 | 0.11 | 2 |
| GPi Neuron | Thalamic Neuron | 1 | 3 | 2 |
| Thalamic Neuron | Cortical Neuron | 1 | 5 | 2 |

where synaptic network connections followed a random connectivity pattern.
